# Accessibility and activity of transcriptional regulatory elements during sea urchin embryogenesis and differentiation

**DOI:** 10.1101/2023.05.14.540718

**Authors:** Cesar Arenas-Mena, Serhat Akin

**Affiliations:** College of Staten Island, and PhD programs in Biology and Biochemistry at the City University of New York (CUNY) Graduate Center.; College of Staten Island, and Biology PhD program at the City University of New York (CUNY) Graduate Center.

## Abstract

Transcriptional regulatory elements (TREs) are the primary nodes of the gene regulatory networks that control development. TREs are identified by PRO-seq and their accessibility by ATAC-seq during sea urchin embryonic development and differentiation. Our analysis identifies surprisingly early accessibility in 4-cell cleavage embryo TREs that is not necessarily followed by subsequent transcription, and an excess of ATAC-seq peaks transcriptionally disengaged during the stages analyzed. Embryonic accessibility shifts are driven by transcriptionally engaged TREs, and PRO-seq transcriptional differences at TREs provide more contrast among embryonic stages than ATAC-seq accessibility differences. TRE accessibility reaches a maximum around the 20-hour late blastula, which coincides with major embryo regionalizations. At the same time, a large number of distal TREs become transcriptionally disengaged, in support of an early Pol II primed model for developmental gene regulation that eventually resolves in transcriptional activation or silencing. A transcriptional potency model based on labile nucleosome TRE occupancy driven by DNA sequences and the prevalence of histone variants is proposed in order to explain the basal accessibility of transcriptionally inactive TREs during early embryogenesis.

**Summary statement:** Genomic profiles deciphering the location and activity of regulatory elements that control gene expression suggest general mechanisms of regulatory potency in early sea urchin embryos.

## Introduction

Developmental gene expression is primarily controlled by transcriptional regulatory elements (**TREs**) that constitute the main nodes of gene regulatory networks (Britten and Davidson, 1969; Peter and Davidson, 2015). High in the regulatory network hierarchy are signaling and transcription factors that determine each other’s expression as well as the expression of downstream differentiation genes at the periphery of the network. These developmental regulators are oftentimes controlled by distal regulatory elements enhancing or silencing the transcription of their “constitutive-like” promoters (Haberle and Lenhard, 2016). In recent models (Andersson et al., 2015), proximal promoters and distal enhancers/silencers are categorized broadly as TREs that use similar mechanisms for the control of distal and local transcription, with productive elongation of protein-coding transcripts representing the outstanding distinctive characteristic of promoters. One of us has proposed an evolutionary causality for these similarities; in short, enhancers/silencers represent the evolutionary cooption of inducible-type promoters for the distal control of originally constitutive regulatory genes in unicellular organisms, which enabled the evolution of complex developmental gene regulation in the lineage leading to metazoans (Arenas-Mena, 2017). Thus, in this scenario enhancer RNAs (eRNAs) represent a property of both proximal and distal TREs rather than evolutionary novelties with a specific regulatory role. In further support of this hypothesis, transcription initiation is necessary for distal enhancer function (Tippens et al., 2020), strengthening the notion of local transcription as an inherent to proximal and distal TREs.

The identification and functional characterization of distal regulatory elements has been greatly facilitated in recent years by diverse genomic profiles and high throughput functional assay methods (Andersson and Sandelin, 2020). Among others, the Assay for Transposase-Accessible Chromatin using Sequencing (**ATAC-seq**) identifies DNA accessibility genome-wide (Buenrostro et al., 2013) while Precision Run-On Sequencing (**PRO-seq**) detects enhancers and promoters transcriptionally engaged (Mahat et al., 2016), that is, TREs with RNA Polymerase II paused or driving productive transcription (Wang et al., 2019). Indeed, using machine learning methods, we found that ATAC-seq and PRO-seq are good predictors of distal enhancer activity empirically validated in sea urchin embryos (Arenas-Mena et al., 2021). It seems that PRO-seq followed by machine learning Discriminative Regulatory Element Detection (**dREG**) does a better job than ATAC-seq to identify the spatial location of TREs validated by unbiased *cis*-regulatory analysis (Arenas-Mena et al., 2021), as expected from its direct report of transcriptionally engaged distal enhancers/silencers and promoters.

Developmental potency requires accessibility to developmental TREs (Andersson et al., 2015; Peter and Davidson, 2015), and the restriction of such potential seems stabilized by differential DNA accessibility among terminally differentiated cells (Arenas-Mena, 2007; Arenas-Mena, 2017; Buenrostro et al., 2018). Thus, developmental potency mechanisms must be driven by the interactions among DNA regulatory sequences, nucleosomes and transcription factors (Brahma and Henikoff, 2020; Michael and Thomä, 2021; Slattery et al., 2014; Zaret, 2020).

Better understanding of the relationships and functional significance of TRE accessibility and transcription has been gained in recent years. TRE accessibility may be stablished first by dedicated “pioneer” transcription factors capable of interacting with nucleosome-bound regulatory DNA, and then recruit chromatin remodeling and histone modifying enzymes that increase local accessibility and facilitate subsequent binding of “settler” transcription factors (Zaret, 2020). Such models assume an antagonistic role of nucleosome occupancy and transcription factor binding, and a default flat DNA accessibility landscape that is disrupted by pioneer recruited enzymatic activities. The relationships between TRE accessibility and transcription are not trivial. Comparative analysis of ATAC-seq and PRO-seq profiles reveals that a substantial number of accessible regions remain transcriptionally disengaged (Arenas-Mena et al., 2021; Wang et al., 2022), favoring models of transcription initiation as a regulated event, as previously discussed (Wang et al., 2022) rather than the unavoidable byproduct of accessibility, as previously proposed (Haberle and Stark, 2018). Furthermore, the formation of the transcription preinitiation complex seems to be associated with greater accessibility than transcription (Wang et al., 2022).

Here we report the analysis of ATAC-seq and PRO-seq profiles during sea urchin embryogenesis and differentiation. The data set provides a comprehensive profile of active TREs that is experimentally validated by prior *cis*-regulatory studies in sea urchin embryos. Large number of distal TREs and the promoters of transcriptionally silent transcription factors are occupied by paused Pol II in early-cleavage embryos, in support of a transcriptionally primed model for developmental gene regulators (Boettiger and Levine, 2009). Surprisingly, our results also reveal accessible but transcriptionally silent TREs in early cleavage embryos, suggesting that a substantial portion of regulatory elements may be accessibility primed by default. The results are integrated in models accounting for the diversity of nucleosome positioning, occupancy and lability that may operate to increase regulatory potency during early embryogenesis.

## Results

### ATAC-seq and PRO-seq profiles during embryonic development and differentiation

In order to understand the developmental changes in TRE accessibility and transcriptional engagement during development and differentiation, we have performed ATAC-seq in 4 cell embryos and ATAC-seq and PRO-seq in 8, 12 and 20 hour embryos, which represent early, mid and late blastula stages, respectively, and in 72 hour larvae and red spherule cells, which represents an adult differentiated state (**Fig. 1**). The ATAC-seq and PRO-seq libraries have good quality metrics and high correlation between biological replicates (**Sup. Fig. 1 A**, **Sup. Fig. 2** and **Sup. Table 1**), with the inherent spread in the lower accessibility ranges (**Fig. 2 A**). For the majority of ATAC-seq and PRO-seq samples (**Sup. Table 1**), we have used our previously published nuclear-prep protocols (Arenas-Mena et al., 2021) with minor variations in adult red spherule cells, see methods. Red spherule cells and 72 hour larvae are more challenging samples, possibly due to endogenous nuclease activity, and we have also tested omni-ATAC (Corces et al., 2017), a method that skips the nuclear prep step (**Fig. 1**). Omni-ATAC in 72 hour larvae was used for subsequent comparative analysis among stages because it results in more correlation among replicates than nuclear prep replicates, not shown. Nevertheless, Omni-ATAC libraries correlate well with nuclear prep libraries in 72 hour larvae and red spherule cells (**Sup. Fig. 1 B**), enabling comparisons with other stages. For all samples, we have used previously adapted PRO-seq methods (Arenas-Mena et al., 2021), and in red spherule cells a quick PRO-seq (**qPRO-seq**) variation (Judd et al., 2020). Library depth is within the same order of magnitude between biological replicates, except for the 8 hour embryos, and among all stages, except for 72 hour larvae and red spherule cells PRO-seq libraries (**Sup. Table 1**), with read coverage below that expected for machine learning dREG TRE predictive saturation (Wang et al., 2019). Nevertheless, the inclusion of PRO-seq data of red spherule cells and 72 hour larvae, while incomplete, provides informative differentiation contrast with the embryonic stages that identify general trends while keeping in mind the expected bias against differentiation dREG TREs derived from the lower coverage in these two differentiated stages, as commented where relevant.

**Figure 1.**
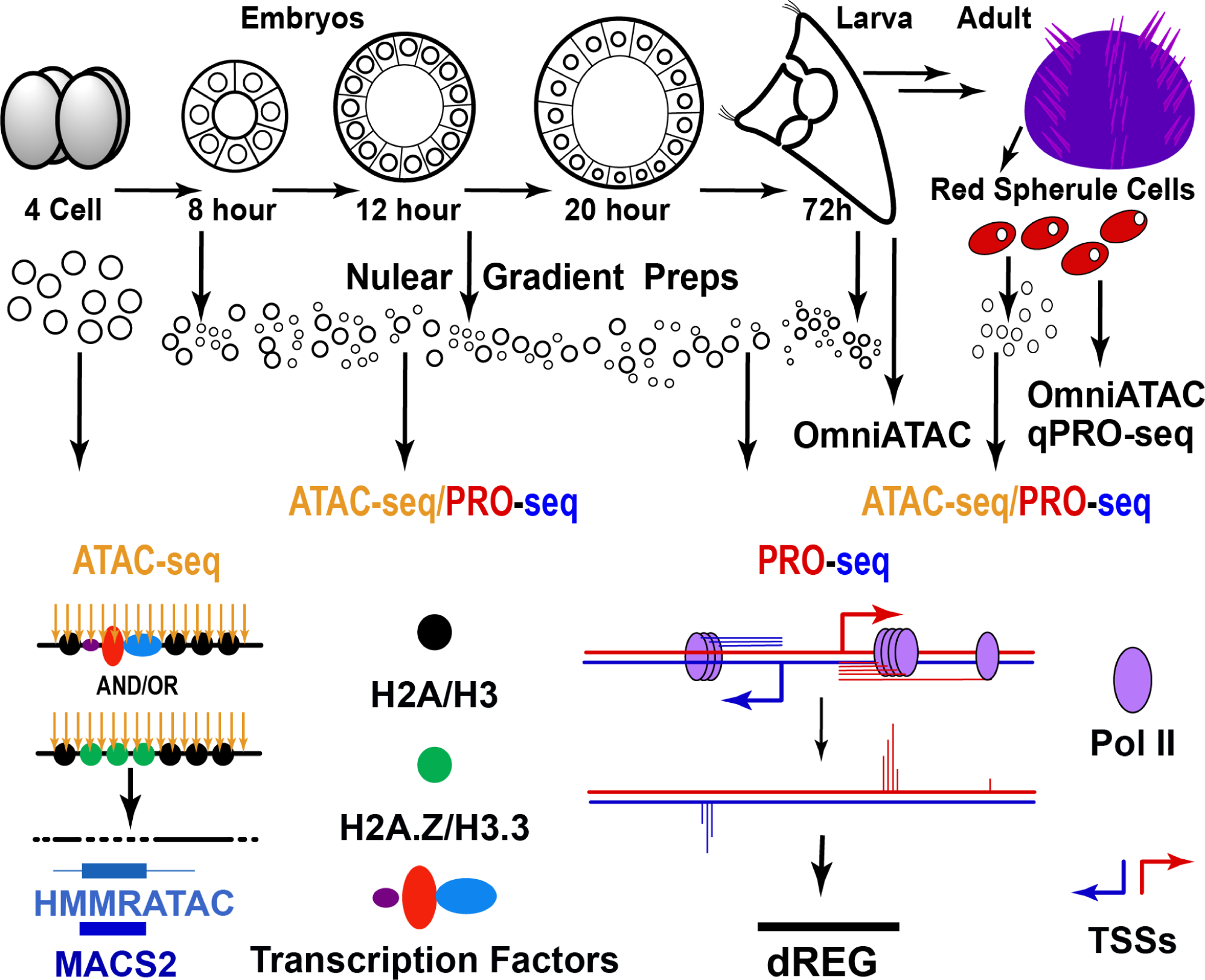
Outline of ATAC- and PRO-seq experiments. Nuclear peps were obtained for all the stages indicated followed by ATAC-seq and PRO-seq experiments. ATAC-seq detects DNA accessibility and PRO-seq the last nucleotide incorporated by transcribing or paused RNA polymerases. Both Omni-ATAC and ATAC-seq libraries have been obtained in 72 hour larva and red spherule cells, with ATAC-seq biological replicate libraries enabling statistical comparisons among stages in adult red spherule cells and omni-ATAC libraries in 72 hour larvae. Quick PRO-seq has only been used in red spherule cells. ATAC-seq peak calls are made by **MACS2** (Model-based Analysis for ChIP-Seq) (Zhang et al., 2008) and **HMMRATAC** (*Hidden Markov ModeleR for ATAC-seq*) (Tarbell and Liu, 2019), with thick lines indicating regions of sub-nucleosomal read length and thin lines flanking nucleosomes. TRE predictions from PRO-seq data are made by **dREG** (*Discriminative Regulatory Element Detection*) (Wang et al., 2019). Pol II, RNA polymerase II; TSSs, transcription start sites; H2A.Z and H3.3 are histone variants.

**Fig. 2.**
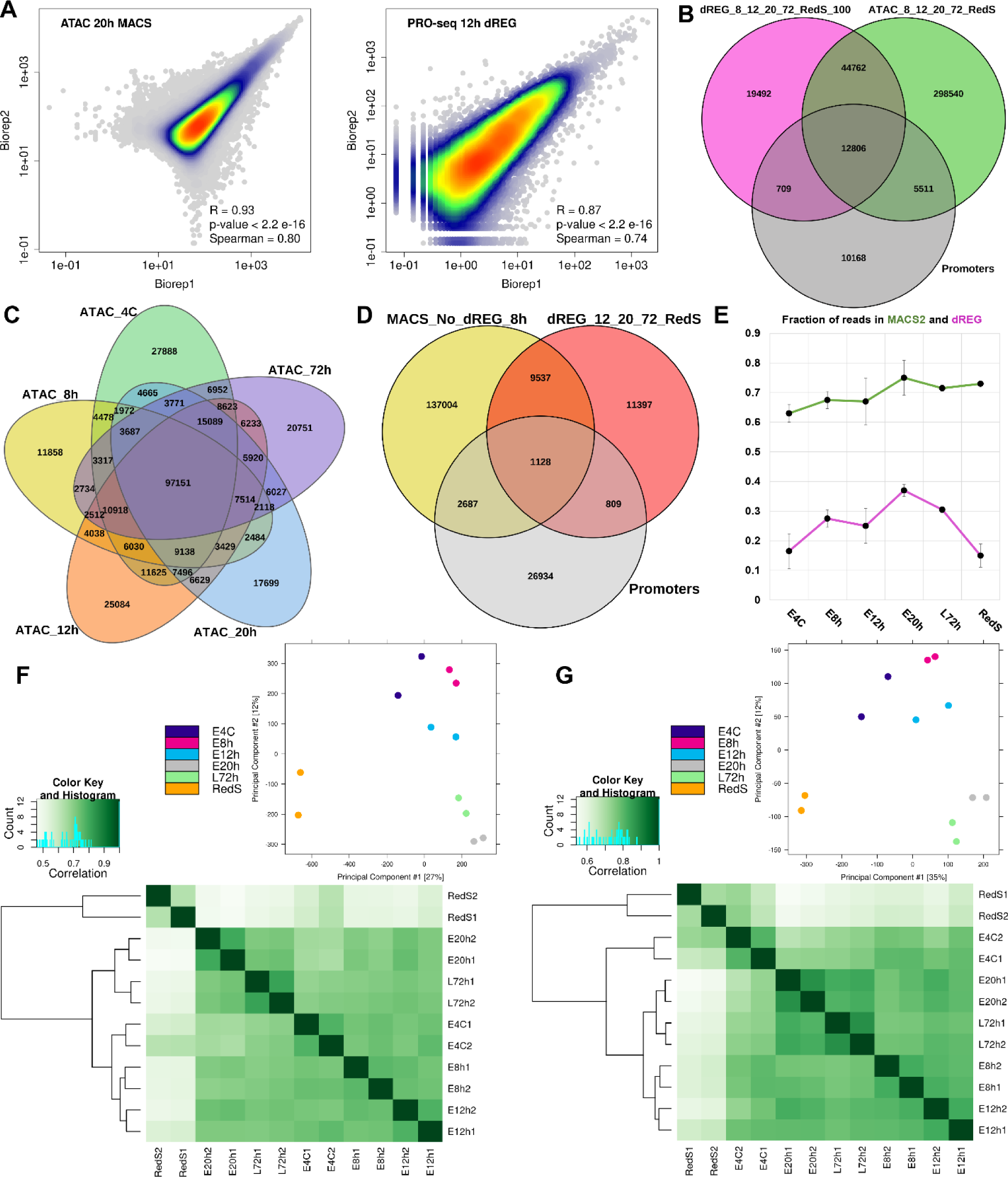
Accessibility correlations between replicates and among stages at MACS2 peak calls and dREG TRE predictions. **A**, plots of ATAC-seq and PRO-seq in reads per million at MACS2 peak calls or dREG TRE predictions of biological replicates for the stages indicated along with reproducibility estimations by Pearson (R) and Spearman correlations. Overlap of points indicated by the color gradient. **B**, overlaps among merged ATAC-seq MACS2 peak calls, PROseq-dREG TRE predictions of embryonic (8, 12, 20 hours), larval (72 hour) and differentiated red spherule cells. **C**. Overlap of MACS2 peak calls among the stages indicated. **D**, overlap among transcriptionally inactive MACS2 peak calls (dREG negative) in 8 hour embryos, promoters and dREG TREs during later stages. **E**, fraction of total reads in biological replicate libraries mapping in merged ATAC-seq MACS2 peak calls and dREG predictions in each stage, error bars for 95 % confidence intervals. **F**, bottom, correlation heatmap using DESeq2 normalized ATAC-seq reads of biological replicates quantified at the combined and merged MACS2 peak calls of all stages. Top, PCA analysis of the same data set. **G**, similar analysis as in F at the combined dREG TRE predictions.

### Incipient TRE accessibility is observed in early embryos prior to TRE activation

Early TRE accessibility across the genome is observed in 4 cell sea urchin embryos, as exemplified by the *Hox11/13b* locus (**Fig. 3**). This result is perhaps unexpected, because few developmental transactions are known to be active at this early stage (Peter and Davidson, 2015), except for the secondary axial specifications that have barely started (Coffman et al., 2009), in agreement with the equal developmental potency of the 4 blastomeres (Reviewed by Horstadius, 1973). Visual inspection of several regulatory loci, not shown, reveals similarly incipient accessibility at developmental TREs that become more accessible and active during much later stages (Barsi and Davidson, 2016; Cui et al., 2017; Nam and Davidson, 2012), when several transcription factors known to drive their enhancer and/or silencer functions are expressed or activated (Howard-Ashby et al., 2006; Tu et al., 2014). For example, in 20 hours, the maximum absolute DNA accessibility of the *Hox11/13b* promoter and module ME (**arrowheads in Fig. 3, left panel**) coincides with maximum *Hox11/13b* expression and module ME enhancer activity (Cui et al., 2017), which experimentally validates the quality of our PRO-seq and ATAC-seq data (**Fig. 3**). However, the module ME and the *Hox11/13b* promoter are already accessible relative to flanking regions in 4 cell embryos (**Double arrowheads, Fig. 3, right panel**), as quantitatively assessed by the underlying peak calls of the Model-based Analysis for ChIP-Seq (MACS2) (Zhang et al., 2008). Incipient accessibility is also observed at TREs of adult red spherule cells (**Triple arrowheads Fig. 3, right panel**). These observations reveal that at least some TREs have above-background accessibility through development and differentiation.

**Figure 3.**
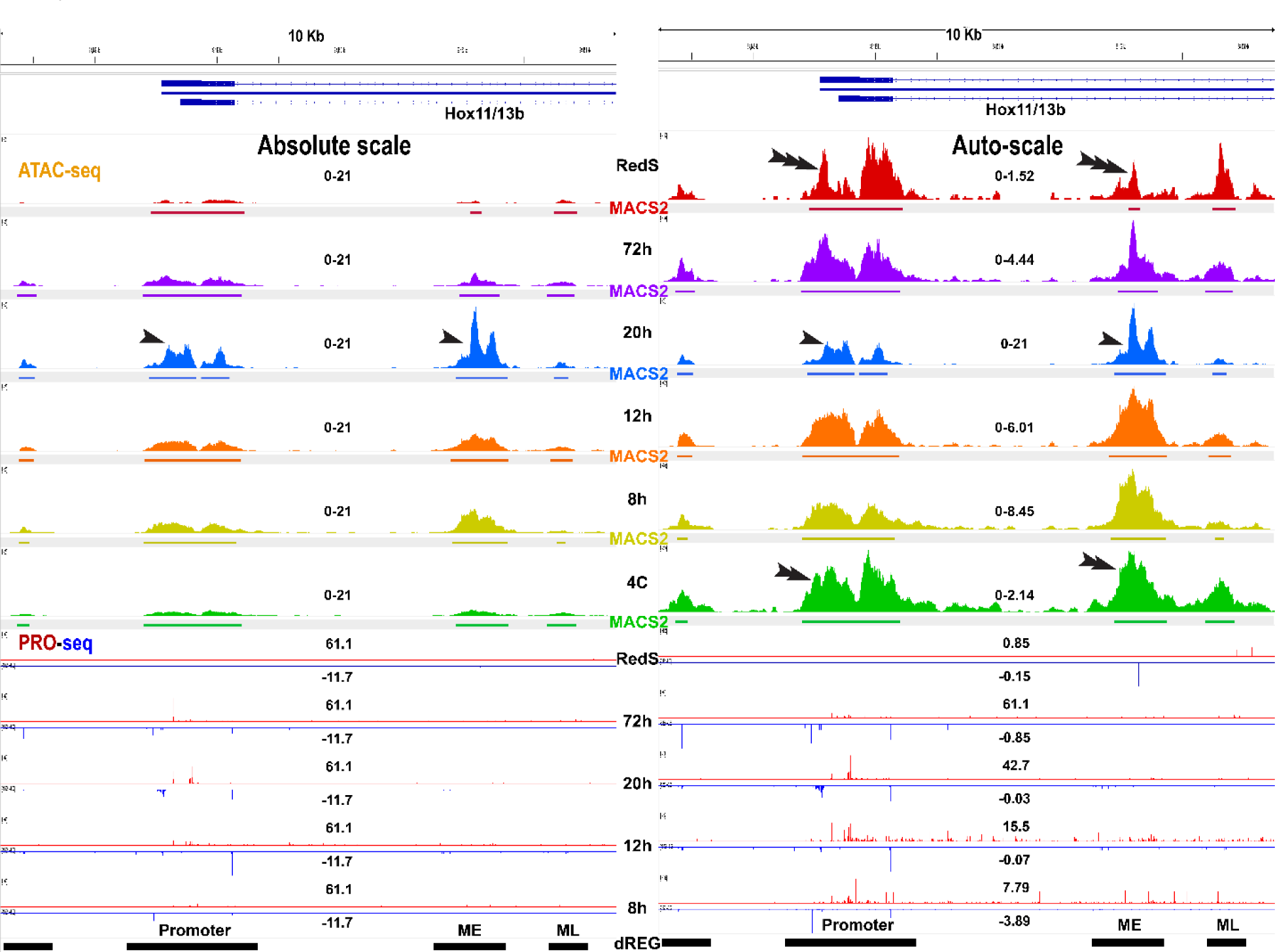
ATAC- and PRO-seq genomic profiles during embryonic development and differentiation at the *Hox11/13b* locus. Genomic profiles in reads per million are the combination of two biological replicates. The PRO-seq run-on transcripts/base is shown for each base in the **plus** (red, top) and **minus** (blue, bottom) strands, with PRO-seq peaks marking the transcriptional pause sites of TSSs. The left panel (**Absolute scale**) shows ATAC-seq profiles set at the maximum scale range of the 20 hour sample in reads per million (**0-21**), and the right panel (**Auto-scale**) with each range set to the maximum value of each track. For PRO-seq the absolute scale is independently set for the plus and minus strands among all samples, with negative values for the minus strand. The dREG TRE predictions are the combined and merged set of the 8, 12, 20, 72 hour and red spherule cells dREG predictions. ME and ML are enhancers whose activity was experimentally tested in reporter constructs (Cui et al., 2017). **4C**, 4 cell embryos; **8-20 h** embryos and **72 h** larva as indicated; **RedS**, differentiated red spherule cells.

Early embryo TRE accessibility does not necessarily anticipate immediate transcriptional activation. The late regulatory module of Hox11/13b (**ML** in **Fig. 3**), only becomes active during postembryonic larval stages (Cui et al., 2017). In addition, significant incipient accessibility is also detected at the *Hox-A7* promoter (**Sup. Fig. 3, single arrowhead**), which remains transcriptionally disengaged, that is, lacks dREG TRE predictions and it is not transcribed. Similarly, there is accessibility without transcriptional engagement in TREs associated with several Hox cluster genes (not shown) that are not transcriptionally active anytime during sea urchin embryonic development but during much later postembryonic larval development, as previously demonstrated (Arenas-Mena et al., 1998; Arenas-Mena et al., 2000), and later confirmed by transcriptome analysis (Tu et al., 2014) and our current PRO-seq data (absence of PRO-seq reads and dREG TRE predictions, as illustrated in **Sup. Fig. 3**). Thus, except for two dREG TREs in 8-hour embryos (**double arrowheads, Sup. Fig. 3**), some of the MACS2 peak calls at the *Hox-A7* locus in 4 cell embryos may correspond to postembryonic distal TREs that are significantly accessible but transcriptionally disengaged during embryogenesis.

The immense majority of accessibility peaks detected during embryonic development and differentiation are not transcriptionally engaged, that is, they do not overlap with dREG TRE predictions (**Fig. 2 B**). This is not surprising because not all the accessibility peaks represent TREs (Klemm et al., 2019), and because additional stages and cell types would have to be sampled in order identify all the TREs in the sea urchin genome, as suggested by the approximately one third of promoters without dREG activity (**Fig. 2 B**). Nearly all transcriptionally engaged promoters are accessible in some stage, but about one third of distal dREG TREs are not (**Fig. 2B**), which suggests that transcription and accessibility are not necessarily coupled, in agreement with the prior report of transcriptionally paused TREs that do not necessarily trigger a substantial increase in accessibility (Wang et al., 2022). The immense majority of accessibility peak calls in embryos, larvae and differentiated red spherule cells overlap among several stages and are already present in 4 cell embryos (**Fig. 2 C** and **Sup. Fig. 4 A**). Accordingly, relatively few accessibility peaks are stage-specific during embryogenesis and larva differentiation (**Fig. 2 C**), with red spherule cells having about twice the number of unique peaks (**Sup. Fig. 4 A**). About half of the dREG TREs active during development or differentiation are already accessible in 4 cell embryos, and the fraction of transcriptionally engaged peaks that overlap accessible peaks increases during the 8 to 12 and 12 to 20 hour embryo transitions (**Sup. Fig. 4 C**). The lower sequencing coverage in 72 hour and red spherule cells should be causing an overall underestimation of the dREG TRE predictions in these two stages and their overlaps cannot be properly evaluated (**Sup. Fig. 4 C**). In 8 hour embryos, where both PRO-seq and ATAC-seq data sets are available, close to half of the TREs that will become active later during embryonic development or in differentiated red spherule cells are already accessible but transcriptionally disengaged, that is, they have MACS2 peak calls that do not overlap dREG peaks in 8 hour embryos (**Fig. 2 D**), and the overlapping proportions among enhancers and promoters are similar. Consequently, transcription is anticipated by accessibility as early as the 8 hour embryo for a large number of TREs. In addition, similar to *Hox-A7* (**Sup. Fig. 3**), more than two thirds of promoters accessible in 8 hour embryos are not transcriptionally engaged in any of the stages analyzed (**Fig. 2 D**). In conclusion, accessibility long anticipates transcriptional activation for a large number of TREs.

### Peak number of TREs with enhanced accessibility is reached in late blastula embryos

In order to identify differential DNA accessibility at accessible peaks (MACS2 peak calls) or transcriptionally engaged TREs (dREG TRE predictions) during development and differentiation, we performed comparative quantitative analyses between stage combinations using background plus edgeR or DESeq2 normalization (**Fig. 4, Sup. Fig. 5 and Sup. Fig. 6**).

Late blastula stage embryos (20 hour) have the highest number of MACS2 and dREG TREs with enhanced accessibility relative to 4 cell embryos (**Fig. 4 B**, and **Sup. Fig. 6 A**). At the same time, there is a substantial number of accessible TREs in 4 cell embryos whose accessibility declines in 20 hour blastula and 72 hour differentiated larvae (**Fig. 4 B**, and **Sup. Fig. 6 A**); although, most of these have intermediate accessibility in 4 cell embryos and few decline to the lowest ranks of blastula and larval stages (**Fig. 4 A** and **Sup. Fig. 5 A**). The differential accessibility analysis concurs with the developmental profile of the fraction of reads in dREG TREs, which reaches a significant maximum in 20 hour embryos (**Fig. 2 E**). A large number of embryonic accessible TREs have lower accessibility in Red spherule cells (**Fig. 4**. and **Sup. Fig. 5 A** and **B**), as expected from a general correlation of declining TRE accessibility during differentiation corresponding with declining regulatory potential and/or general transcriptional activity, while a relatively small number of TREs have enhanced accessibility, possibly associated with differentiation functions in this cell type. According to both EdgeR and DESeq2, the sharpest increase in accessible TREs happens during the 12 to 20 hour blastula transition (**Sup. Fig. 6 D**), which associates with a declining rate of cleavage and longer transcriptionally active interphases, while the sharpest decrease in accessible TREs happens during the transition to differentiation from the 20 hour blastula to 72 hour larva (**Sup. Fig. 6 D**).

**Figure 4.**
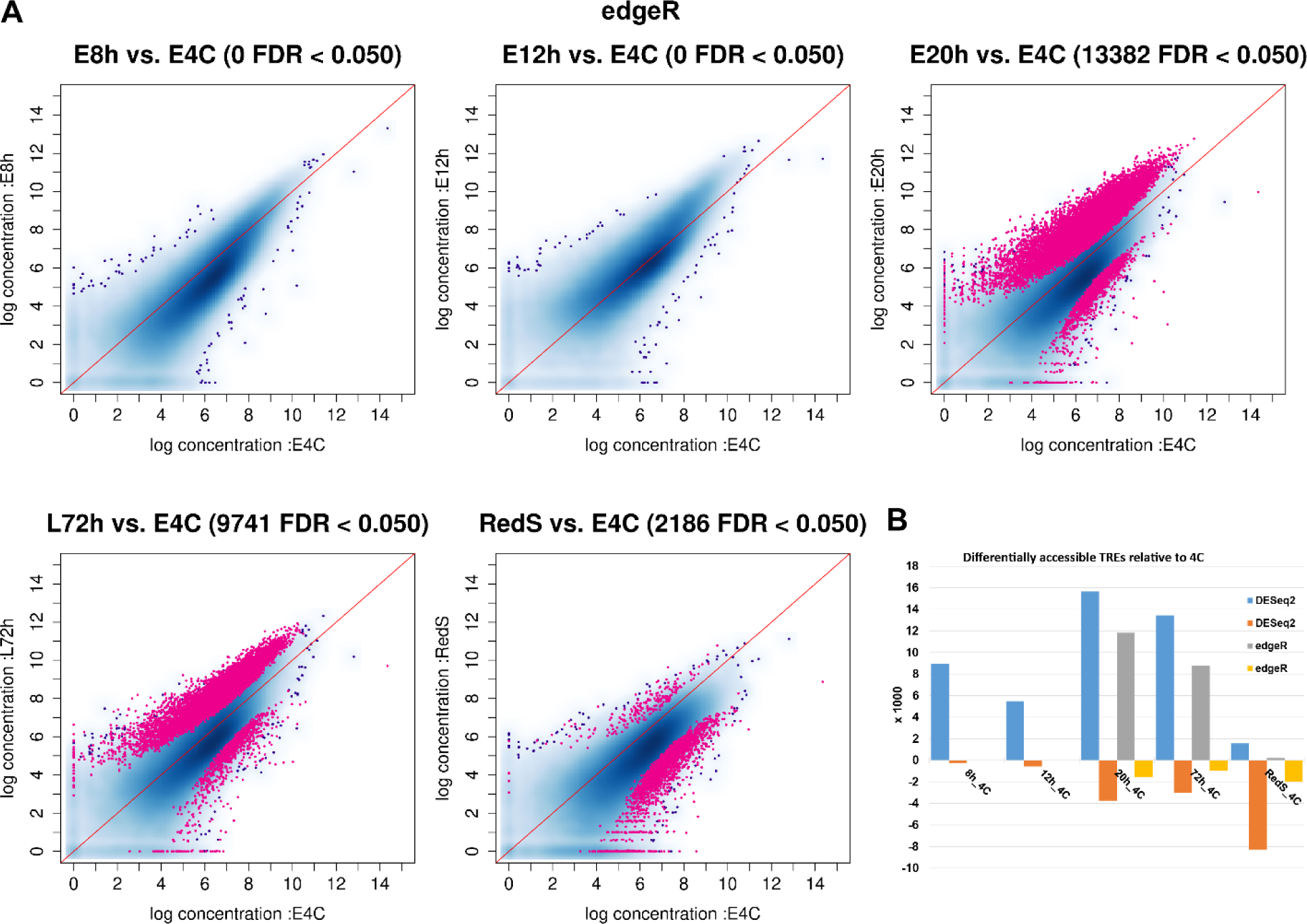
Differential ATAC-seq reads at dREG TREs during development and differentiation. **A**, plot of background plus edgeR normalized ATAC-seq read counts of 8, 12 and 20 hour embryo, 72 hour larva and red spherule cells against 4 cell embryos at the merged dREG TREs of 8, 12 and 20 hour embryos, 72 hour larva and Red Spherule cells (**RedS**) . In pink, TREs with significantly higher or lower counts at false discovery rate (FDR) < 0.05. **B**, summary of the number of the combined set of dREG TRE predictions with accessibility significantly above or below the 4 cell embryo stage after background plus DESeq2 or edgeR normalization. **RedS**, red spherule cells.

These general trends are consistent with the two normalization methods used, background plus edgeR and background plus DESeq2, see methods. However, edgeR normalization seems more stringent and detects no significant accessibility differences between 4 cell embryos and 8h and 12h embryos (**Fig. 4**), while DESeq2 detects more up and down regulated dREG predicted TREs in all stages (**Sup. Fig. 5**). The distinct effects of both normalization methods on the fold changes across the accessibility range and on the false discovery rates over fold changes is illustrated by comparison of MA and volcano plots, respectively, between 8 hour and 20 hour embryos (**Sup. Fig. 5 C**). Given the uncertainties of functionally tested TRE ground truth, we consider appropriate to report the results using both normalization methods.

### Differential accessibility primarily involves transcriptionally engaged TREs

Accessibility at dREG predicted TREs is significantly variable during development (**Fig. 2 E**, pink line), unlike most accessibility differences between stages for the MACS2 peak calls (**Fig. 2 E,** green line). Indeed, high correlations are found between DESeq2 normalized reads at MACS2 peak calls for embryonic (4C, 8h, 12h, 20h) and larval stages (72h) (**Fig. 2 F**). Experimental reproducibility is validated by the grouping between biological replicates and by the secondary affiliation according to sequential developmental stages, with red spherule cells (RedS) occupying their expected outgroup position. However, particularly for accessibility correlations at MACS2 peak calls (**Fig. 2 F, bottom**), dendrogram branch lengths between replicates and those among embryonic and larval stages are similar, meaning that overall accessibility differences have similar ranges within and between stages. Similar results are obtained when the ATAC-seq reads are computed at the merged set of dREG predicted TREs in 8h, 12h, 20h, 72h and RedS, except for the 4 cell embryos, which are more different to the subsequent embryonic stages and occupy now a basal position (**Fig. 2G**), in agreement with the global accessibility roughly estimated by the fraction of reads in dREG TREs (**Fig. 2 E**). For both the MACS2 peak calls and dREG predicted TREs, almost identical results to those of DESeq2 (**Fig. 2 F** and **G**) are obtained when the analysis is performed after edgeR or library depth normalization (not shown). The correlation results summarized by the histograms (**Fig. 2 F and G, bottom**) are confirmed by principal component analyses of the same data sets (**Fig. 2 F and G, top**). All together, these results suggest that the developmental accessibility differences among stages primarily corresponds to transcriptionally engaged TREs.

Indeed, the majority of the MASCS2 peak calls that gain significant accessibility during embryonic development and larva differentiation become transcriptionally engaged sometime during development and differentiation, as demonstrated by the similar number of dREG TREs and MACS2 peak calls with higher accessibility relative to 4 cell embryos (**Sup. Fig. 6 A**), which primarily correspond to MACS2 peak calls overlapping dREG predictions (Compare **Sup. Fig. 6 A** and **B**). Thus, transcriptional engagement of TREs corresponds with increasing accessibility during development, as anticipated by the increasing overlap of MACS2 and dREG (**Sup. Fig. 4 C**). Conversely, the great majority of the MASCS2 peak calls that significantly loose accessibility relative to 4 cell embryos during embryonic development and larva differentiation remain transcriptionally disengaged, that is, they do not overlap with the merged set of dREG TRE predictions (Compare **Sup. Fig. 6 A** and **C**). A slightly different result is found in red spherule cells, where dREG overlapping TREs represent a small fraction of those with higher accessibility (Compare **Sup. Fig. 6 A, B and C**). However, this can be explained in part by the expected underrepresentation of red spherule dREG TREs due to the lower PRO-seq sequencing depth in this cell type mentioned before, which will undermine the identification of red spherule cell specific dREG TREs that are nevertheless accessible.

The identity of the accessibility shifts correspond to biological functions known to be activated during development, including those related to general metabolic and regulatory functions, as exemplified by GO term analysis of the gene promoters overlapping dREG TREs with differential accessibility (**Sup. File 1**).

### Differential PRO-seq signal at dREG TREs during the early to late blastula transition

We performed a similar differential PRO-seq analysis with the suitable profiles (**Fig. 5** and **Sup. Table 1**) at the same combined dREG TREs used in the differential ATAC-seq analysis (**Fig. 2**). The PRO-seq samples for 72 hour larvae and red spherule cells (**Sup. Table 1**) have been excluded from the quantitative evaluation of the number of differentially Pol II engaged dREG TREs (**Fig. 5 B**), because they could be biased due to their relatively lower number of reads (**Sup. Table 1**). Interestingly, for a substantial number of dREG TREs, the PRO-seq signal declines during the 8 to 20 hour blastula transition, while it increases for a relatively small fraction of TREs (**Fig. 5 A** and **B**). Similar to the ATAC-seq results, edgeR is more stringent than DESEQ2 (**Fig. 5 B**), the data is validated by the grouping according to stage (**Fig. 5 C** and **D**), except for a one step deviation of a 72 hour biological replicate (**Fig. 5 C**). Overall, the correlations among TRE transcriptional engagement estimations by PRO-seq (**Fig. 5 C**) are significantly lower (**Fig. 5 E**) than those among TRE accessibility estimations by ATAC-seq (**Fig. 2 G**). In other words, PRO-seq offers clearly more contrast among developmental stages than ATAC-seq.

**Figure 5.**
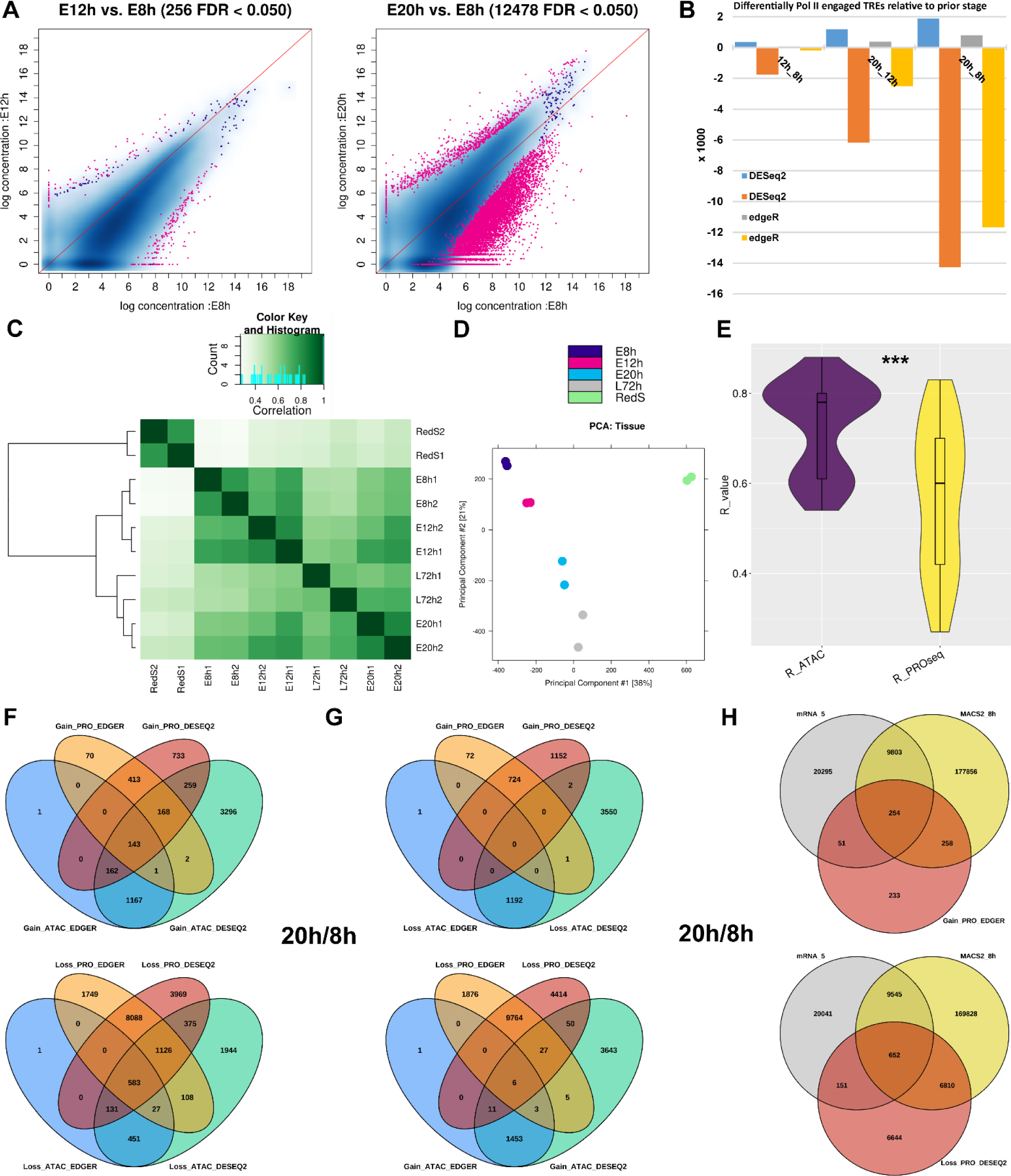
Differential PRO-seq reads at dREG TREs during embryonic development. **A**, representative plots of edgeR normalized PRO-seq read counts of 8 and 20 hour embryos relative to 8 hour embryos at the merged dREG TREs of 8, 12 and 20 hour embryo, 72 hour larva and red spherule cells. In pink, TREs with significantly higher or lower counts at false discovery rate (FDR) < 0.05. **B**, summary of the number of the combined set of dREG TRE predictions with PRO-seq reads significantly above or below the embryo stage in the second term, such as 20 hour relative to 12 hour embryo in 20h_12h. **C**, correlation heatmap using edgeR normalized PRO-seq reads of biological replicates quantified at the combined and merged peak calls of all stages. **D**, PCA analysis of the same data set in C. **E**, comparison of the ATAC-seq and PRO-seq Pearson correlation coefficients in C and Fig. 2 G, Wilcox text p-value = 3.036 x 10^-6^, **\*\*\***. **F** and **G**, intersect among dREG TREs with differential gain and loss of PRO-seq or ATAC-seq signals between 20 to 8 hour embryos using edgeR or DESEQ2 normalization as indicated. **H**, intersect of the dREG TREs with differential PRO- and ATAC-seq signals in F with the 5’ side of mRNAs.

Analysis of differential ATAC-seq and PRO-seq signals at dREG TREs during the 8 to 20 hour embryo transition identifies overlapping and non-overlapping sets for both upregulated and downregulated signals (**Fig. 5 F**). ATAC-seq identifies more upregulated TREs (**Fig. 5 F**, **top**), while PRO-seq identifies a larger number of TREs that loose signal (**Fig. 5 F**, **bottom**). Nevertheless, there is very little overlap between up and down regulated PRO-seq and ATAC-seq TREs (**Fig. 5 G**), revealing that the differences are primarily due to the detection of distinct sets rather than a negative correlation between ATAC-seq and PRO-seq signals, as expected. EdgeR and DESEQ2 identify complementary and overlapping TREs for PRO-seq signals, while edgeR ATAC-seq TREs almost entirely represent a subset of those identified by DESEQ2 (**Fig. 5 F**).

The immense majority of the PRO-seq downregulated TREs are distal enhancers or silencers, while half of the upregulated TREs are promoters (**Fig. 5 H**). This suggests a general transcriptional decommissioning of distal TREs during the early to late blastula transition. A more balanced proportion of promoters to enhancers/silencers is found between ATAC-seq up and downregulated TREs, although with substantially different ratios depending on the normalization method (not shown). There is general agreement among PRO-seq, ATAC-seq and mRNA profiles for 8 to 20 hour PRO-seq upregulated promoters, as illustrated for developmental transcription factors *Dlx*, *GATAe*, *Tbx2*, and *NK2.2* (**Fig. 6**). However, for the downregulated counterparts, while there is parallel decline in PRO-seq, ATAC-seq and mRNA profiles for some promoters, as exemplified for Zf-835, the PRO-seq signal decline in other promoters does not clearly correspond to a parallel decline in ATAC-seq signal and associates with genes that although transcriptionally primed are not expressed in 8 and 20 hour embryos, such as *Sox9* and *Dbx1* (**Sup. Fig. 7**). All of the above suggest that TRE accessibility and transcription changes are not necessarily correlated.

**Figure 6.**
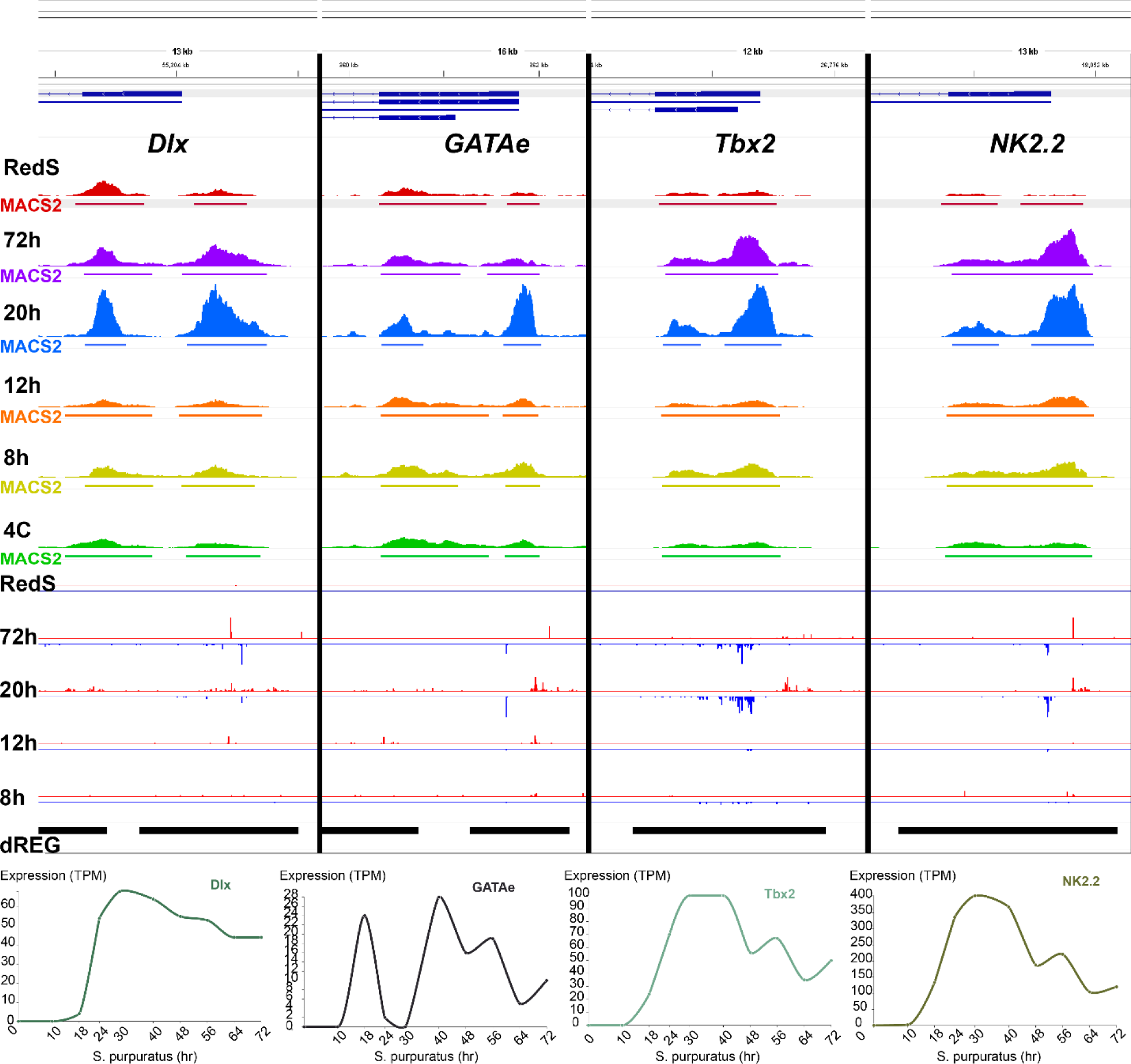
ATAC-seq, PRO-seq and mRNA expression profiles of 8h to 20 hour PRO-seq upregulated promoters. TPM, transcripts per million.

### Most TREs maintain basal accessibility in both early embryos and differentiated cells

The immense majority of dREG TREs that are differentially accessible relative to 4 cell embryos in 20 hour embryos, 72 hour larvae and red spherule cells are already accessible in 4 cell embryos, that is, they are detected by MACS2 (**Sup. Fig. 8 A**). This is the case not only for TREs that gain accessibility but also for those that lose it, which represent the majority of differentially accessible TREs relative to 4 cell embryos in red spherule cells (**Sup. Fig. 8 A**).

These results agree with the visual inspection of accessibility profiles in early embryos (exemplified in **Fig. 3** and **Sup. Fig. 3**). Given the extensive overlap of accessibility peaks in red spherule cells with dREG TREs and accessibility peaks in embryos and larvae (**Sup. Fig. 4 A and B**), we asked if differentially accessible dREG TREs in early embryos maintain basal accessibility during terminal differentiation, or if such extensive overlap could only be explained by constitutively accessible regions. We found that, despite the prevalent suppression of embryonic dREG TRE accessibility in red spherule cells (**Sup. Fig. 6 E**), the majority of differentially accessible dREG TREs in 20 hour embryos and 72 hour larvae that are accessible in 4 cell embryos (**Sup. Fig. 8 A**) remain detectable by MACS2 in red spherule cells (**Sup. Fig. 8 B**). These results reveal that the basal TRE accessibility observed in 4 cell embryos is also maintained for most TREs after differentiation.

Basal TRE accessibility could depend on transcription factor binding or “labile nucleosomes”, which have been proposed to be associated with A/T rich sequences (Chereji et al., 2019). However, our analysis reveals significantly lower A/T content at ATAC-seq accessible regions, dREG TREs and promoters (**Sup. Fig. 9 A**), as illustrated by several TREs (**Sup. Fig. 9 B**). We performed an additional accessibility analysis using an ATAC-seq dedicated method: Hidden Markov ModeleR for ATAC-seq (HMMRATAC) (Tarbell and Liu, 2019)(**Sup. Fig. 9 B**). The analysis generally coincides with MACS2, although HMMRATAC peak calls are generally broader (**Sup. Fig. 9 B**). In addition to accessibility, HMMRATAC identifies the location of flanking nucleosomes (**Fig. 1**), thin lines in **Sup. Fig. 9 B.** However, it cannot distinguish between nucleosome free regions and labile nucleosomes at the center of accessible peaks, because both associate with sub-nucleosomal read lengths (Chereji et al., 2019). Visual analysis reveals A/T sequence depletion at the central accessible regions relative to flanking nucleosomes (**Sup. Fig. 9 B)**, in agreement with A/T sequence distributions of the narrower MACS2 and dREG TRE predictions (**Sup. Fig. 9 A**). Furthermore, there is no correlation between ATAC-seq and A/T content at accessible regions (MACS2 positive) that are transcriptionally disengaged (dREG negative) in 8 hour embryos (**Sup. Fig. 9 C**). This reveals that A/T content does not explain the accessibility variation of inactive early sea urchin embryo TREs.

## Discussion

Our results reveal intriguingly early and generalized TRE accessibility in the pluripotent blastomeres of the 4 cell sea urchin embryo, and in 8 hour embryos we demonstrate TRE accessibility long before transcriptional activation (**Fig.2 D**, **Fig. 3 and Sup. Fig. 3**). TRE activity is also anticipated by earlier accessibility in mammalian embryos (Thurman et al., 2012; Wu et al., 2016), stem cells (Meers et al., 2019), oocytes and sperm (Jung et al., 2019), and spatial decoupling of accessibility and transcription has been reported at inactive regulatory elements in Drosophila and sea urchins embryos (Calderon et al., 2022; Shashikant et al., 2018). Therefore, at least during embryogenesis and perhaps also during differentiation, there are basal levels of TRE accessibility that do not concur in time or space with TRE transcription. Altogether, our results are against models of transcription as the unavoidable corollary of accessibility (Felsenfeld, 2014), and favor TRE transcription as a process regulated at several limiting steps that may include increased DNA accessibility (Wang et al., 2022). Therefore, differential PRO-seq data analysis better approximates the distinct transcriptional states deployed during development (**Fig. 5 E**), as expected from its direct report of transcriptional engagement.

Our results also demonstrate that differential accessibility during embryogenesis is primarily associated with dREG TREs (**Sup. Fig. 6**), therefore, other structural and replicative processes triggering accessibility (Klemm et al., 2019; Wang et al., 2022) are expected to be less prevalent or to have more constant accessibility. The majority of the accessible peaks are not transcriptionally engaged during development (**Fig. 2 D**), but many of these accessible regions possibly have transcriptional roles during postembryonic stages yet to be characterized, as previously exemplified (**Sup. Fig. 3**); and this is possibly the case because a substantial number of accessibility differences are detected in differentiated adult red spherule cells (**Sup. Fig. 6 E**) and a substantial number of promoters remain transcriptionally disengaged (dREG negative) and inaccessible among the stages analyzed (**Fig. 2 B**). Therefore, if differential transcriptional engagement remains the main source of differential accessibility during postembryonic development and differentiation, then a large number of accessibility peaks detected during embryogenesis may actually correspond to transcriptionally engaged TREs during postembryonic stages.

It remains to be determined if TRE accessibility in early embryos is nucleosome-bound or nucleosome-free. Early embryo transcriptional potency may be stablished by dedicated “pioneer” transcription factors that can interact with nucleosome-bound regulatory DNA, and then recruit chromatin remodeling and histone modifying enzymes that increase local accessibility and facilitate subsequent binding of “settler” transcription factors (Zaret, 2020) (**Fig. 7, flat landscape model**). In zebrafish embryos, transcription factor binding sites for Pou5f3, SoxB1, and Nanog are covered by transposase-accessible nucleosomes prior to the sequence-specific binding of Pou5f3 and Nanog, which increases accessibility during the main wave of zygotic genome activation (Veil et al., 2019). The authors proposed that the apparently contradicting overlap of early accessibility and nucleosome occupancy may be due to partial nucleosome occupancy, the fraction of cells in a sample with a nucleosome at a particular loci. However, high DNA-accessibility and nucleosome occupancy are not mutually exclusive and can overlap at “labile nucleosomes” that could explain the increased accessibility at inactive TREs (**Fig. 2 D** and **Sup. Fig. 3**), as elegantly demonstrated by quantitative micrococcal nuclease methods that account for nucleosome release and destruction kinetics (Chereji et al., 2019). So, nucleosome lability could explain early embryo TRE accessibility, yet our results (**Sup. Fig. 9**) are against the proposed model of simple A/T sequence enrichment determining nucleosome lability (Chereji et al., 2019). Nevertheless, DNA-sequence-driven nucleosome lability at inactive TREs prior to transcription factor binding remains possible, because empirical models of nucleosome occupancy demonstrate that A/T sequence periodicities rather than overall GC content determine the low intrinsic DNA bending expected at labile nucleosomes and nucleosome depleted regions (Basu et al., 2021). Therefore, we propose that some TREs may be inherently primed with increased accessibility directly set by intrinsic DNA sequences determining the strength of DNA-histone interactions (**Fig. 7, sequence-primed TRE model**). Of course, the flat landscape and sequence-primed models and their variations are not exclusive and are expected to occur at different TREs across the genome.

**Figure 7.**
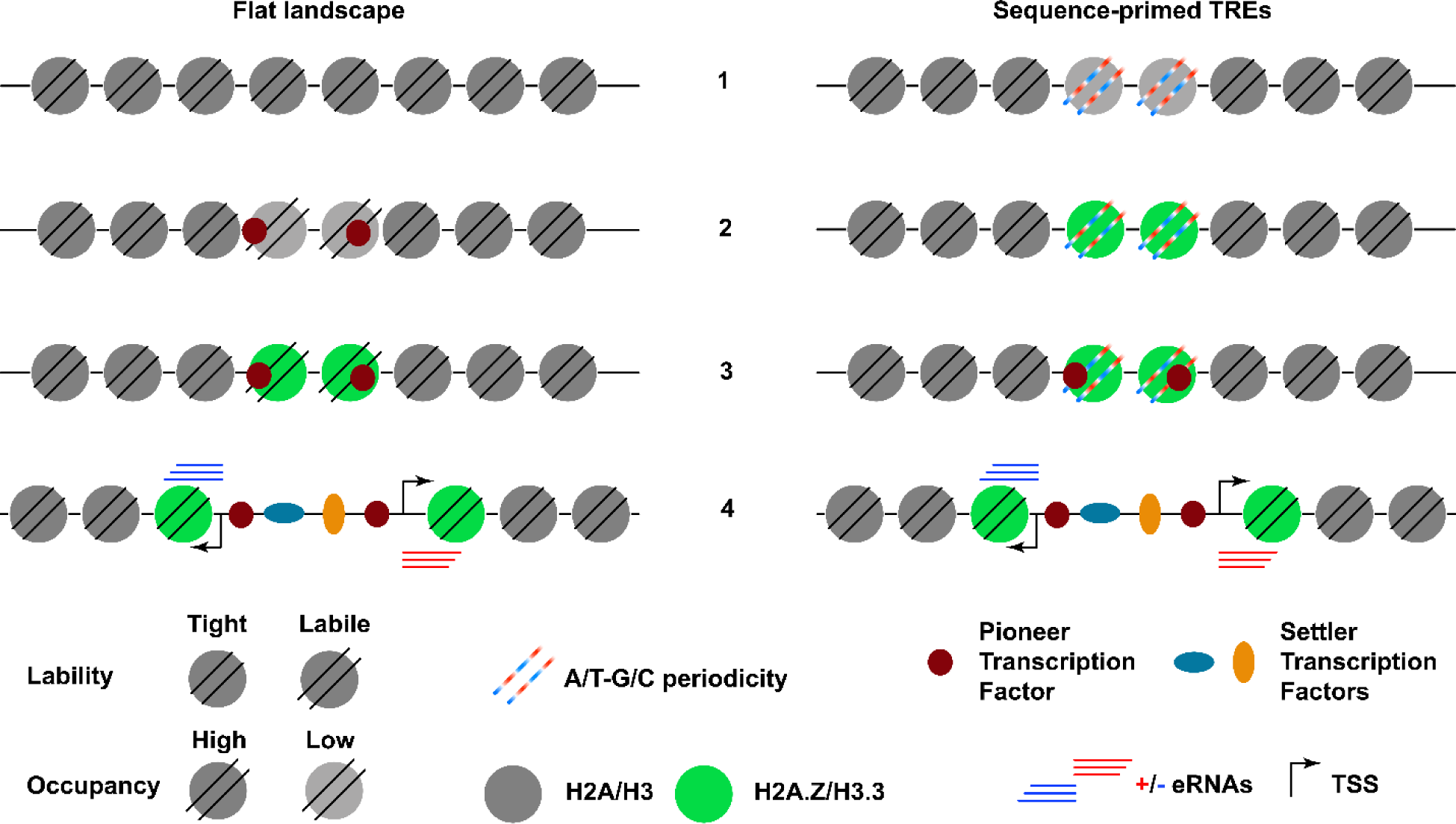
Generic sequence of events, 1 to 4, leading to TRE activation in flat accessibility landscape (left) or sequence-primed TREs (right). **Left**, early during development, TREs are “hidden” in a generally **flat** chromatin accessibility **landscape**, **1**; zygotic pioneer transcription factors recruit chromatin enzymatic activities that loosen DNA-histone interactions, lower nucleosome occupancy, **2**, and replace canonical replication histones with H2A.Z and H3.3 histone variants, **3**, which further increase DNA accessibility and facilitate settler transcription factor binding and enable TRE transcriptional activation, **4**. **Right**, in **sequence-primed TREs**, regulatory elements inherently have incipient accessibility thanks to A/T-G/C sequence periodicities that weaken DNA interactions with canonical histones H2A and H3 and cause labile nucleosomes, **1**, that are replaced by highly abundant histone variants H3.3 and H2A.Z, **2**, and facilitate pioneer transcription factor activity, **3**, eventually leading to TRE activation, **4**, or stable silencing (not shown). Alternative chromatin modifications, other than remodeling or histone variants, and/or sequence of events (**2 to 3**) would fit both models, which are non-exclusive and primarily defined by their initial state, **1**. **eRNAs**, enhancer RNAs (or upstream and coding RNAs in promoters); **TSS**, transcription start site.

The highest number of more accessible TREs in 20 hour embryos coincides with maximum levels of histone variant *H2A.Z* mRNA expression (Ernst et al., 1987; Tu et al., 2014) with the gene-expression echinoid “hourglass phylotypic stage” (Malik et al., 2017), and with the deployment of crucial developmental gene regulatory networks (Li et al., 2014; Oliveri et al., 2008; Peter and Davidson, 2011). H2A.Z is non-essential for individual cell survival but high levels are required during sea urchin embryogenesis (Hajdu et al., 2016), as in other embryos. Nucleosomes with the H2A.Z/H3.3 histone variant combination are labile, have high turnover rates (Henikoff and Smith, 2015; Jin et al., 2009; Soboleva et al., 2014), and locate at regulatory regions previously thought to be nucleosome free due to their artefactual depletion during high-salt nucleosome preparations (Jin et al., 2009). H2A.Z is associated with partially unwrapped nucleosomes (Brahma and Henikoff, 2020) and modulates the stability and mobility of nucleosomes (Rudnizky et al., 2016). Furthermore, our previous analysis found that the expression of H2A.Z during sea urchin and polychaete embryonic and postembryonic development clearly correlates with developmental potency and against differentiation (Arenas-Mena et al., 2007), suggesting a general transcriptional potency role that possibly involves partnership with H3.3 (Arenas-Mena, 2007). This hypothesis has gained strength after the demonstrated requirement of H2A.Z to maintain the pluripotent state of stem cells (Hu et al., 2013) and the requirement of H3.3 during nuclear transfer reprogramming (Jullien et al., 2012). Thus, it could be that maximum transcriptional regulatory potential may be reached at the mesenchyme blastula stage and not prior, which would explain why trans-fating experiments that rely on relatively late ectopic transcription factor gene expression easily work during embryogenesis (Oliveri et al., 2008; Peter and Davidson, 2015).

Interestingly, our analysis also identifies a large number of distal TREs whose PRO-seq signal is suppressed during embryonic development, and a substantial number of them do not overlap accessibility peaks (**Fig. 5 A**, **F** and **H**). This favors a transcriptional rather than an accessibility primed model for distal enhancers during development. In addition to transcriptionally active sites, PRO-seq signals are also associated with H3K4me3 and H3K27ac bivalent sites primed in embryonic stem cells for subsequent activation or repression (Wang et al., 2022). Therefore, our results favor the previously proposed Pol II “pre-loaded” model (Boettiger and Levine, 2009) that would posit widespread distal TRE transcriptional engagement during early embryogenesis. This model may be particularly relevant to developmental gene regulation, which strongly dependents on distal TREs (Haberle and Stark, 2018).

## Methods

Adult purple sea urchins, *Strongylocentrotus purpuratus*, were obtained from The Cultured Abalone Farm, Goleta, California, and maintained in a closed aquarium system under a carrot diet. Embryos were obtained as previously described (Hajdu et al., 2016).

Nuclear preps, ATAC-seq and PRO-seq in embryos and larvae followed previously stablished protocols (Arenas-Mena et al., 2021), except that for red spherule cells the lysis buffer during the nuclear prep was supplemented with 0.1 % NP40, 0.1% Tween-20, and 1% BSA, and the density gradient solutions were supplemented with 1% BSA to diminish nuclear shearing. Red spherule cells were purified from the coelomic fluid of adult sea urchins following a previously stablished method (Smith *et al*., 2019). OmniATAC in 72 hour embryos raised at 15 °C and red spherule cells was performed as previously described (Corces *et al*., 2017).

Quick PRO-seq in red spherule cells was performed according to a previously stablished method (Judd et al., 2019) with minor modifications. Permeabilization and wash buffers were supplemented with 1% BSA, cells were collected in a fixed angle centrifuge at 0 °C and 1,000 G for 10 minutes. The run-on reaction was done at 15 °C for 15 minutes. Dynabeads Myone ® Streptoavidin T1 beads were used to pull down the biotinylated RNAs. Free biotinylated nucleotides were removed by filtration with Sigmaspin^TM^ post-reaction clean-up columns. On-Bead 5’ Decapping reaction was done using NEBuffer 2 buffer (NEB, B7002S ®). Off-Bead Reverse Transcription followed the instructions in SuperScript ® IV Reverse transcriptase kit. The Quick PRO-seq PRO-seq libraries were purified from 5% acrylamide gel to remove adaptor dimers.

ATAC-seq and PRO-seq reads were mapped to the *Strongylocentrotus purpuratus* Sp5.0 genome and MACS2 peak calls and dREG TRE predictions were also performed as previously described (Arenas-Mena et al., 2021), except that the version 2.1.1 of the ENCOCE-DCC atac-seq-pipeline (ENCODE ATAC-seq pipeline. https://github.com/ENCODE-DCC/atac-seq-pipeline) with default parameters was used and dREG TRE predictions are extended 100 bp in order to better match the data distribution (Wang et al., 2019).

Quality control biological replicate plots (**Fig. 2 A**) were performed as previously described (Arenas-Mena et al., 2021). Venn diagrams were elaborated with ChiPpeakAnno (Zhu et al., 2010), with “connected peaks” set to “minimum”. IGV (Thorvaldsdottir et al., 2013) was used for all genome browser visualizations with bigwig files normalized to reads per million. Gene expression profiles (**Fig**. **6**) were from (Tu et al., 2014) and graphics obtained from Echinobase (Arshinoff et al., 2022).

Differential gene expression analysis and visualization was facilitated by the DiffBind package (Ross-Innes et al., 2012). DESEQ2 and EdgeR normalizations were performed using default parameters plus background normalization was used for ATAC-seq. For comparative analysis, raw PRO-seq 3’ end read counts were computed at the merged set of dREG TRE predictions for each biological replicate using the bigWig package https://github.com/andrelmartins/bigWig). No background normalization was combined with DESEQ2 and EdgeR normalization in the PRO-seq differential data analysis.

Promoters were defined by the +/-150 bp region flanking the first nucleotide of all transcripts. The BEDTools (Quinlan and Hall, 2010) nuc function was used to compute AT content of different genomic profiles (**Sup. Fig. 9 A**) and at 10 bp windows across the genome (**Sup. Fig. 9 B**).

## Acknowledgements

We would like to thank Charles Danko and his lab members for general guidance prior to the elaboration of this manuscript and for providing access to the Cornell BioHPC cloud computing. Thanks to Professor Zhong Wang to upgrade dREG for the analysis of the Sea Urchin Genome. Thanks to the BioHPC cloud support personnel for assistance installing and resolving computational issues. Thanks to Rory Stark for the upgrade of DiffBind to easily accept PRO-seq data. Thanks to the support personnel at CSI.

## Data Sharing

The datasets generated and analyzed during the current study are available at NCBI GEO under accession number ####### (pending).

---

**Supplementary Table 1.** Total and biological replicate number of reads of ATAC-seq, OmniATAC, PRO-seq and Quick PRO-seq replicates. FRiP, fraction of reads in peaks.

| Tissue | Total reads | Replicate | Reads | FRiP | MACS2 | Method |
| --- | --- | --- | --- | --- | --- | --- |
| E4C | 76600668 | E4C1 | 14977476 | 0.18 | 177672 | ATAC-seq |
|  |  | E4C2 | 61623192 | 0.15 | 300000 | ATAC-seq |
| E8h | 38829176 | E8h1 | 15037464 | 0.29 | 166453 | ATAC-seq |
|  |  | E8h2 | 23791712 | 0.26 | 219134 | ATAC-seq |
| E12h | 71699574 | E12h1 | 35016184 | 0.21 | 281462 | ATAC-seq |
|  |  | E12h2 | 36683390 | 0.29 | 263477 | ATAC-seq |
| E20h | 51650236 | E20h1 | 28815064 | 0.4 | 210179 | ATAC-seq |
|  |  | E20h2 | 22835172 | 0.34 | 207684 | ATAC-seq |
| L72h | 57935034 | L72h1 | 24655132 | 0.31 | 221367 | OmniATAC |
|  |  | L72h2 | 33279902 | 0.3 | 245445 | OmniATAC |
| RedS | 46687530 | RedS1 | 20508416 | 0.15 | 255398 | ATAC-seq |
|  |  | RedS2 | 26179114 | 0.15 | 272233 | ATAC-seq |

**Supplementary Table 1.** Total and biological replicate number of reads of ATAC-seq, 4 OmniATAC, PRO-seq and Quick PRO-seq replicates.
| Tissue | Total reads | Replicate | Reads | Method |
| --- | --- | --- | --- | --- |
| E8h | 48168387 | E8h1 | 3541161 | PRO-seq |
|  |  | E8h2 | 44627226 | PRO-seq |
| E12h | 26860872 | E12h1 | 12978762 | PRO-seq |
|  |  | E12h2 | 13882110 | PRO-seq |
| E20h | 51946709 | E20h1 | 42529842 | PRO-seq |
|  |  | E20h2 | 9416867 | PRO-seq |
| L72h | 3995491 | L72h1 | 2244502 | PRO-seq |
|  |  | L72h2 | 1750989 | PRO-seq |
| RedS | 11349313 | RedS1 | 6639747 | qPRO-seq |
|  |  | RedS2 | 4709566 | qPRO-seq |

**Supplementary Figure 1.**
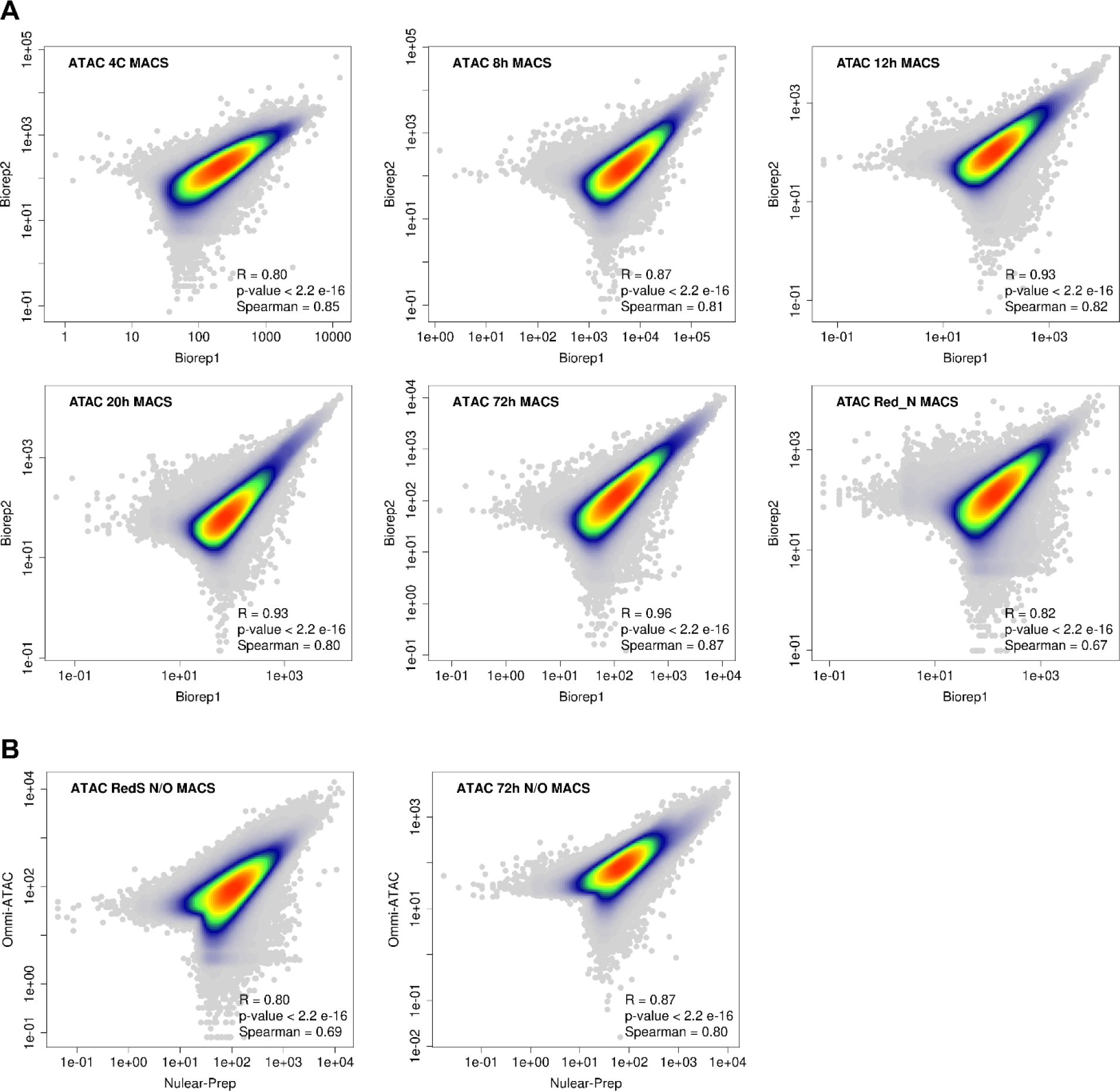
Correlation between biological replicates and ATAC-seq methods. **A,** plot of reads per million at MACS2 peak calls of biological replicates for the stages indicated along with reproducibility estimations by Pearson (R) and Spearman correlations. Overlap of points indicated by the color gradient. **B,** plot as in A for ATAC-seq (nuclear prep) and omni-ATAC in red spherule cells and 72 hour larvae.

**Supplementary Figure 2.**
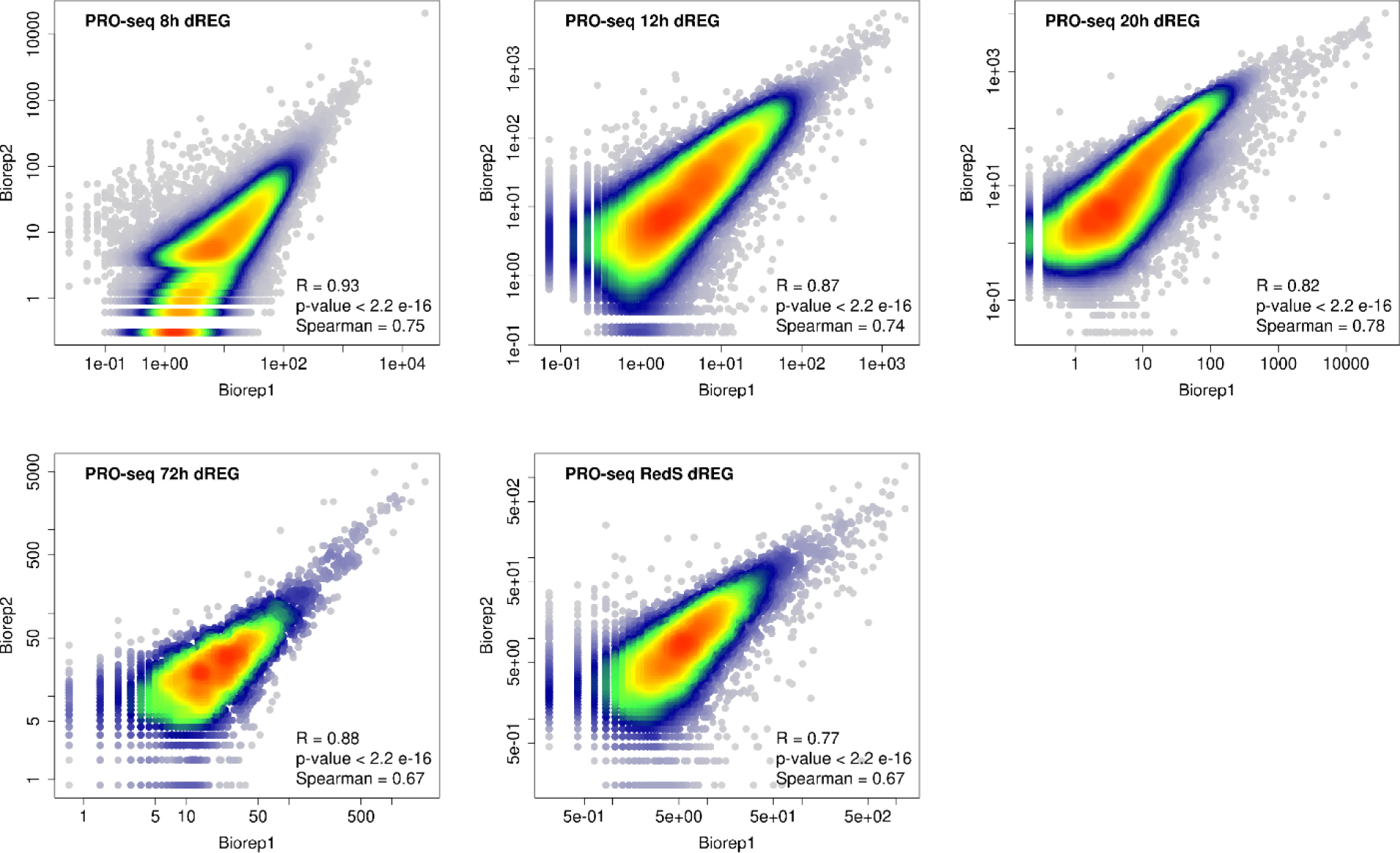
Correlation between PRO-seq biological replicates. Plot of reads per million at dREG peak calls of biological replicates for the stages indicated along with reproducibility estimations by Pearson and Spearman correlations. Overlap of points indicated by the color gradient. The number of mapped reads is 12 times higher between replicates in 8 hour embryos (Sup File 2), which may explain the distortion in the lowest signal ranks, despite the strong overall correlation.

**Supplementary Figure 3.**
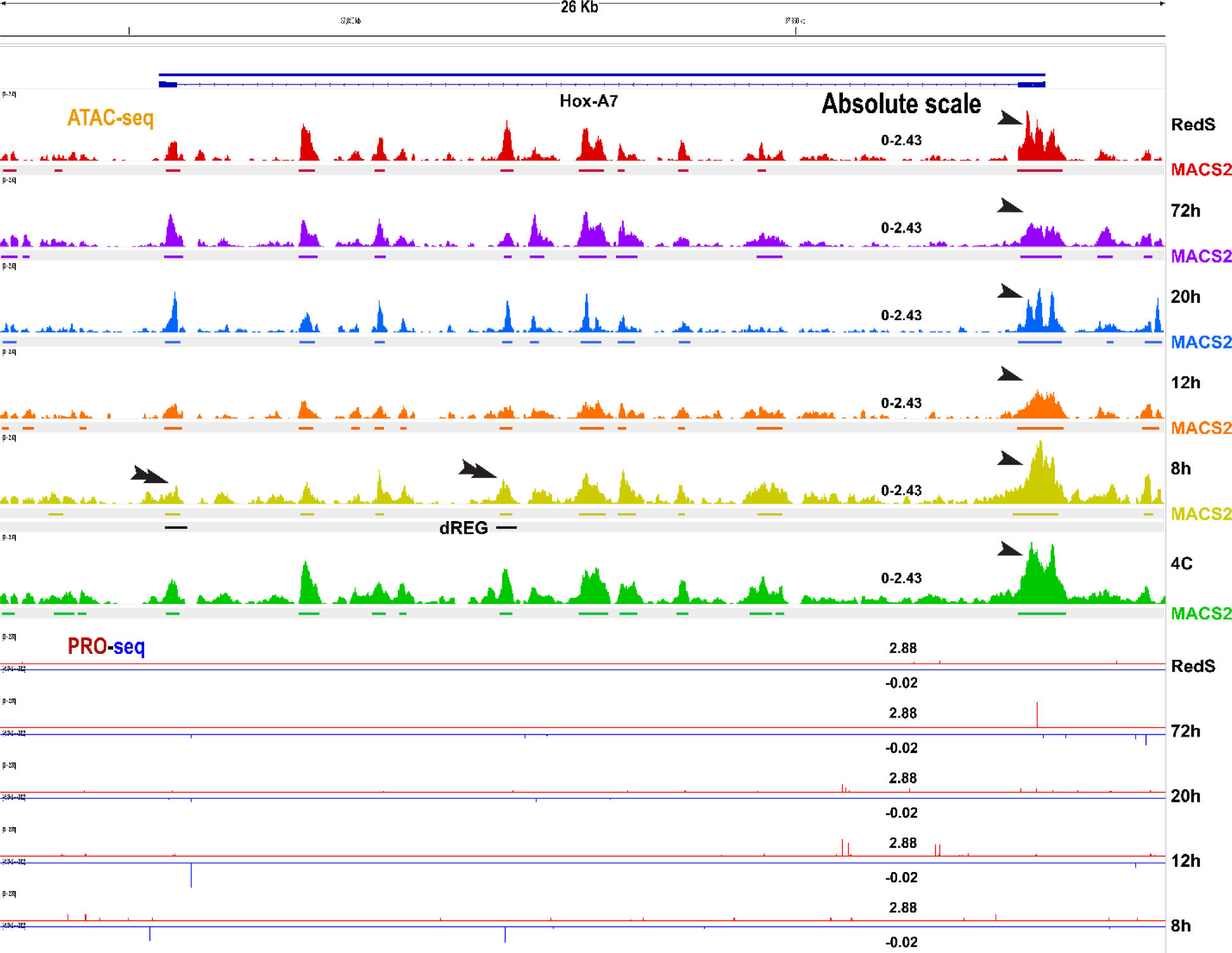
Early embryo TRE accessibility does not necessarily anticipate subsequent transcription. The *HoxA7* promoter is accessible but transcriptionally inactive during embryogenesis and larva differentiation, which corresponds with the lack of PRO-seq signals and dREG peak calls in 12 and 20 hour embryos and 72 hour larvae, but it is accessible and MACS2 positive in 4 Cell embryos (**arrowhead**). Only two dREG TREs are detected in 8 hour embryos (**double arrowheads**).

**Supplementary Figure 4.**
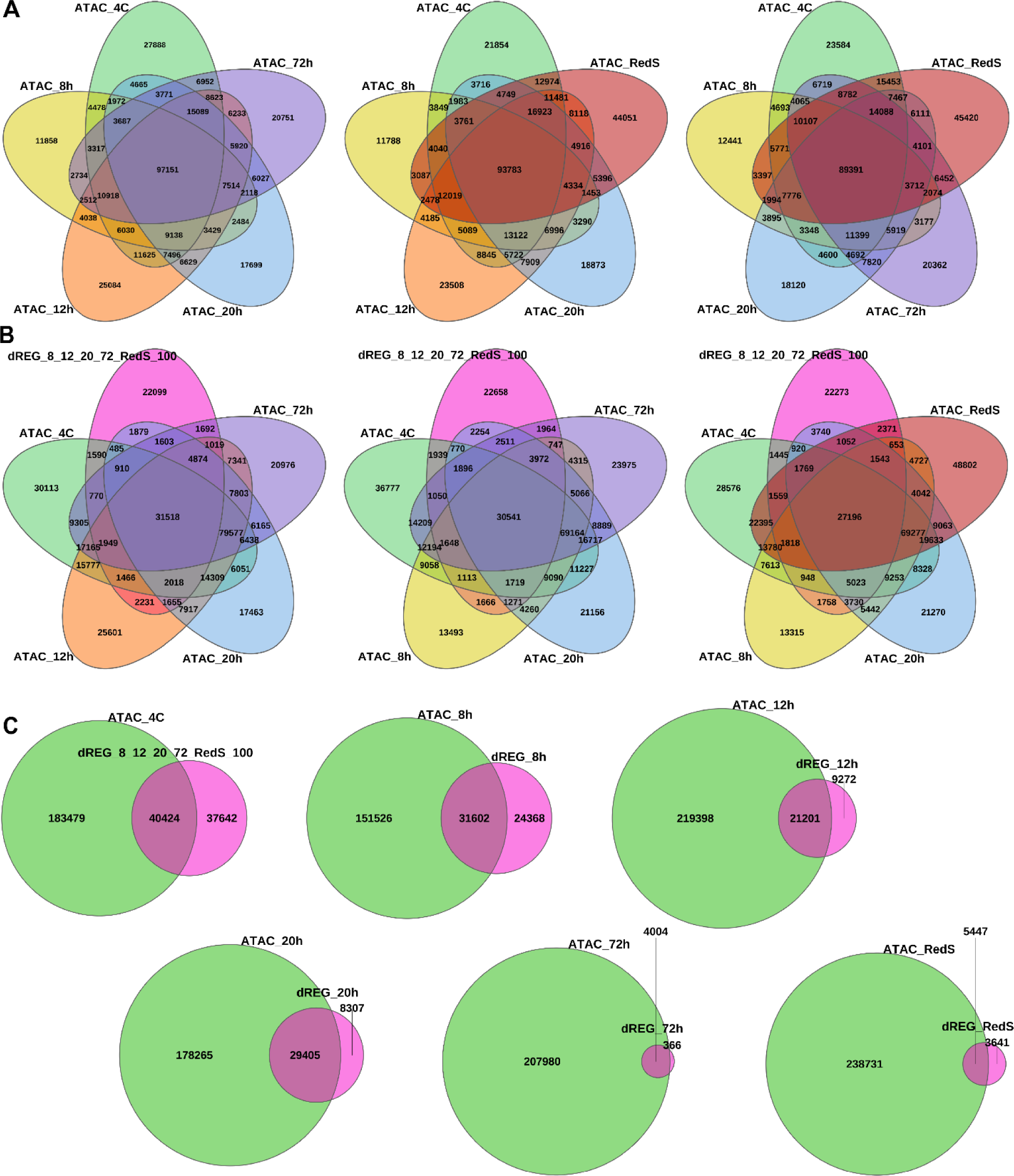
Overlap among and between MACS2 and dREG TRE predictions. **A** Overlap of MACS2 peak calls among the stages indicated. **B**, overlap between ATAC-seq MACS2 peak calls and PROseq-dREG TRE predictions of merged or individual stages as indicated. **C**, between ATAC-seq MACS2 peak calls for different stages and PROseq-dREG TRE predictions of merged or individual stages as indicated.

**Supplementary Figure 5.**
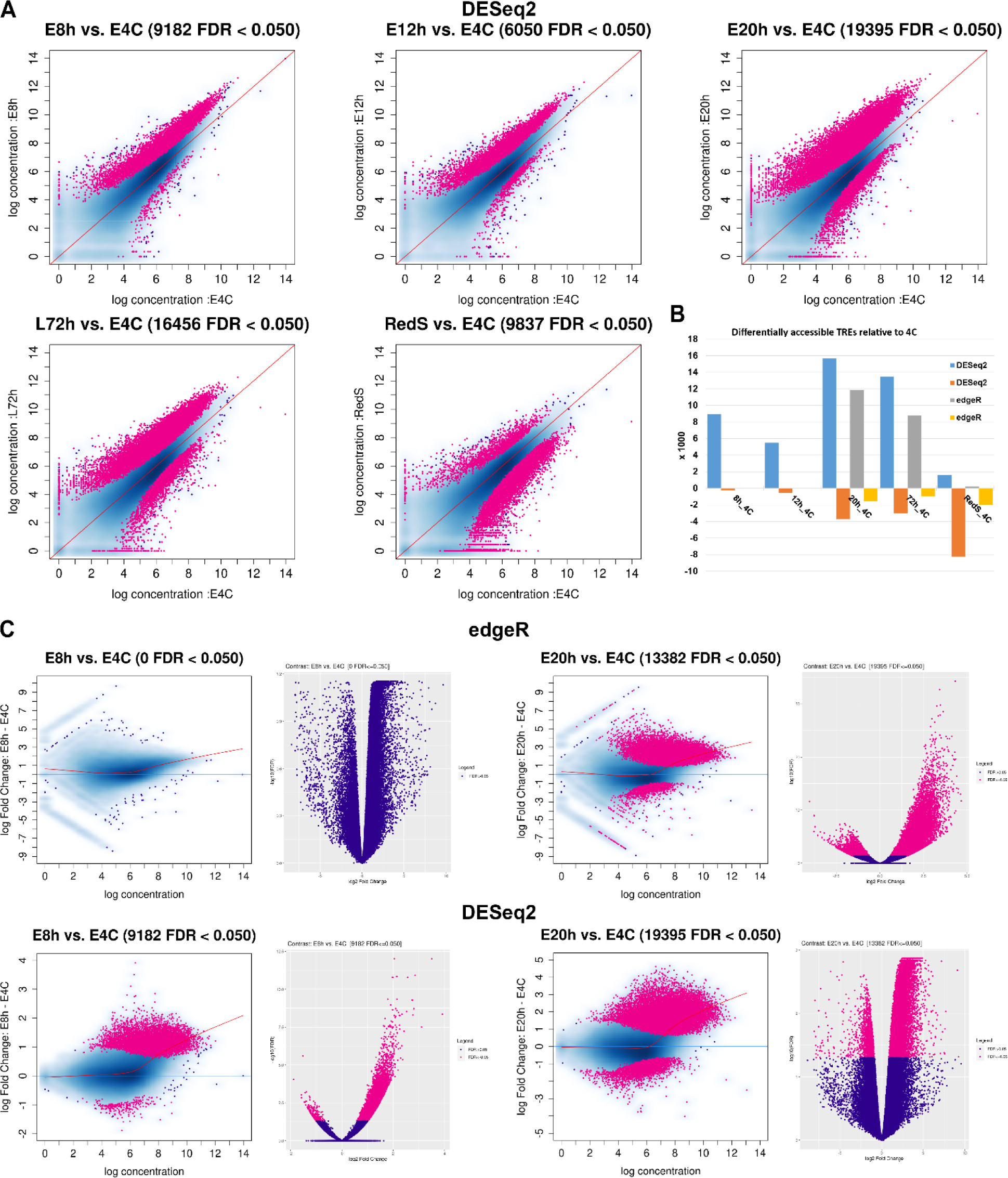
Differential TRE accessibility during development and differentiation. **A**, plot of background plus DESeq2 normalized merged-dREG-TRE read counts of 8, 12 and 20 hour embryo, 72 hour larva and Red Spherule cells (RedS) against 4 cell embryo merged-dREG-TRE read counts. In pink, TREs with significantly higher or lower counts at false discovery rate (FDR) < 0.05. **B**, summary of the number of the combined set of dREG TRE predictions with accessibility significantly above or below the 4 cell embryo stage after background plus DESeq2 or edgeR normalization. **C**, volcano and MA plots of 8 hour and 20 hour data sets shown in A and Fig. 4.

**Supplementary Figure 6.**
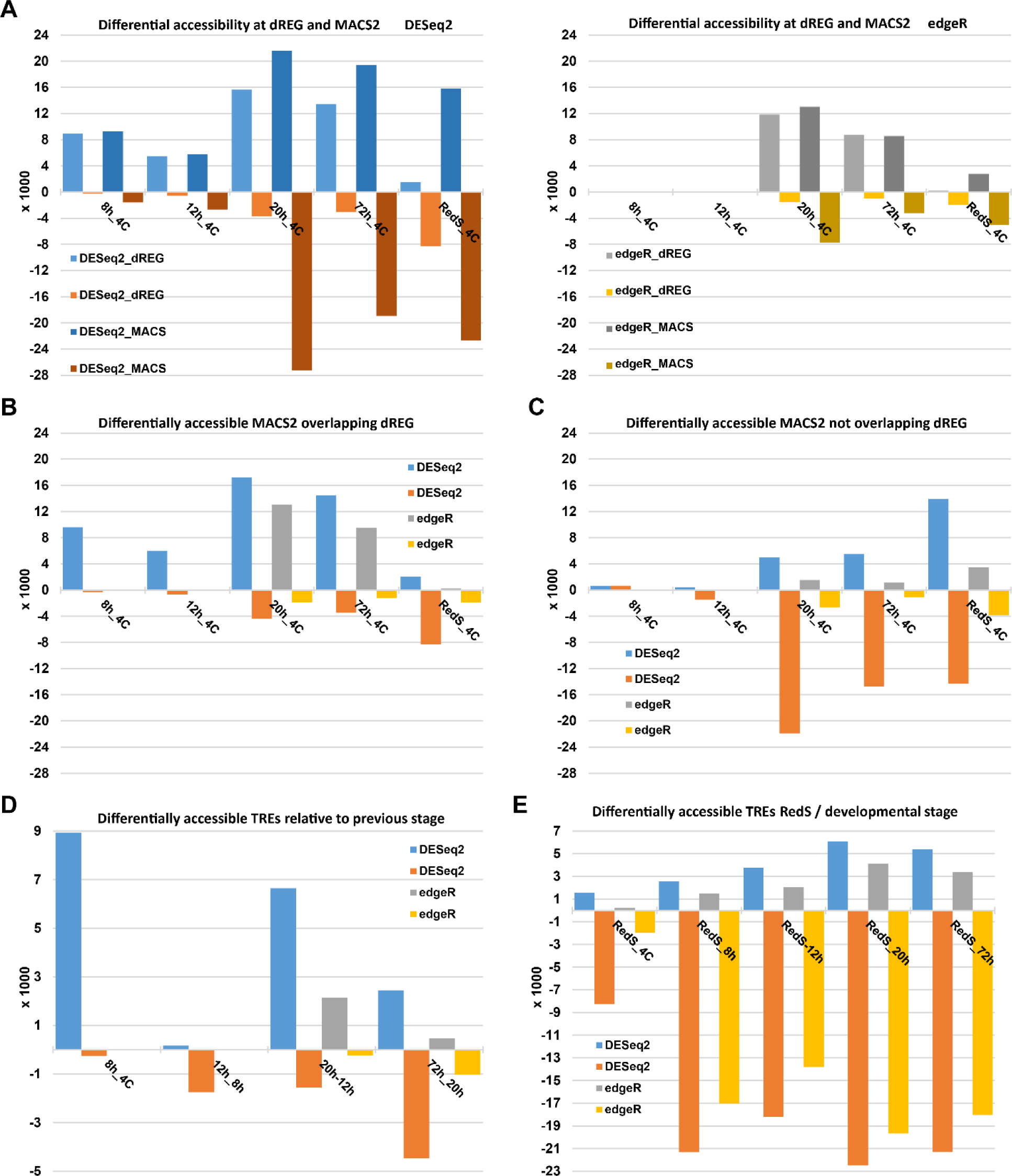
Differential TRE accessibility in MACS2 and dREG TRE predictions. **A**, number of combined stage MACS2 peak calls or dREG TRE predictions with accessibility significantly above or below the 4 cell embryo stage after background plus DESeq2 (left) or edgeR (right) normalization. **B**, number of combined stage MACS2 peak calls overlapping dREG TRE predictions with accessibility significantly above or below the 4 cell embryo stage after background plus DESeq2 or edgeR normalization. **C**, number of combined set of MACS2 peak calls not overlapping dREG TRE predictions with accessibility significantly above or below the 4 cell embryo stage after indicated normalization. **D,** Differential accessibility at dREG TRE predictions relative to previous stage after indicated normalization. **E**, Differential accessibility at dREG TRE predictions relative to differentiated red spherule cells.

**Supplementary Figure 7.**
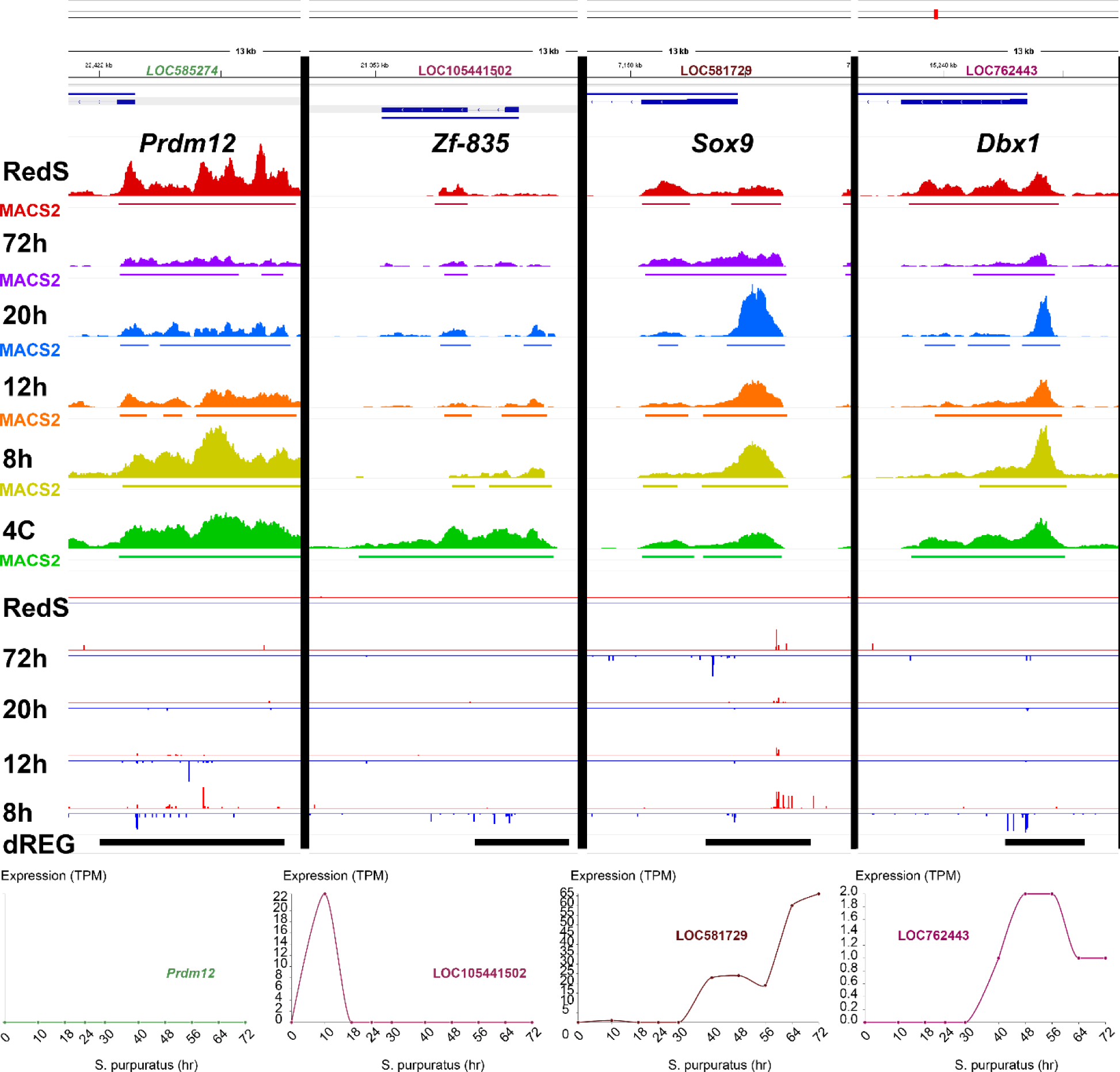
ATAC-seq, PRO-seq and mRNA expression profiles of 8 to 20 hour PRO-seq downregulated promoters. TPM, transcripts per million.

**Sup. Figure 8.**
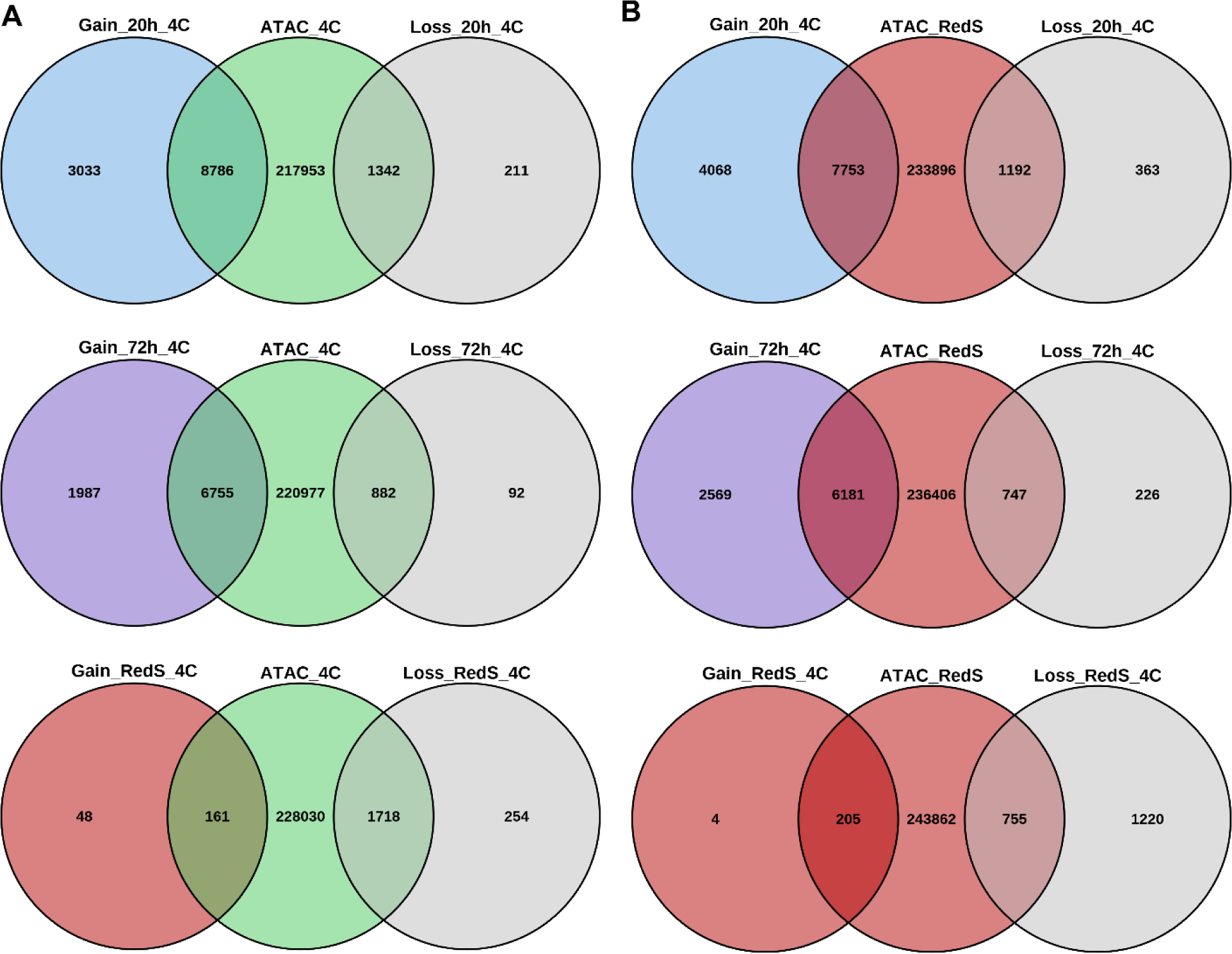
Overlap of differentially accessible dREG TREs with accessible regions in early embryos and differentiated red spherule cells. **A,** Overlap of MACS2 peaks in 4 cell embryos with combined dREG TREs found to be edgeR differentially accessible in 20 hour embryos, 72 hour larvae and red spherule cells relative to 4 cell embryos (Fig. 4). **B,** overlap of MACS2 peak calls in red spherule cells with the same set of dREG TREs in A.

**Supplementary Figure 9.**
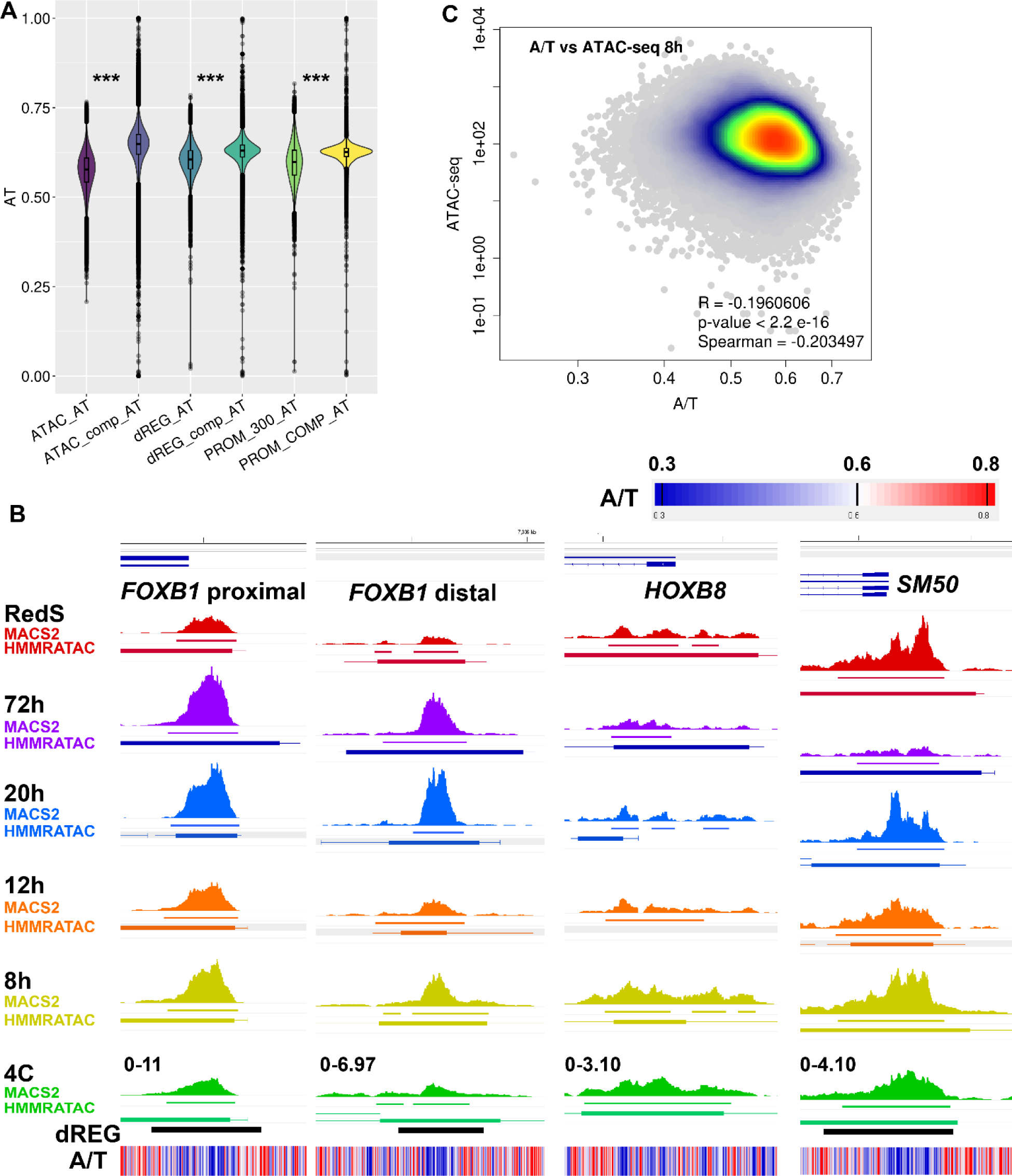
Analysis of A/T content accessibility dependencies. **A,** comparison of A/T content at ATAC-seq, dREG TRE predictions and promoters with their complementary genomic regions; ***, Wilcox text p-value < 2.2 x 10^-16^. Promoters are defined as the 300 bp region centered at the annotated 5’ end of transcripts. **B**, A/T fraction at regions overlapping MACS2, dREG and/or HMRRATAC peak calls, with thick lines for nucleosome free regions and thin lines for flanking nucleosomes. Color scale centered at the A/T genome average of ∼ 0.63. Absolute scale range in reads per million indicated in the 4C track. **C**, plot of 8 hour embryo ATAC-seq signal at MACS2 peak calls that lack dREG TRE peak calls against A/T content illustrative sample of A/T content at several loci.

